# Morphology and synapse topography optimize linear encoding of synapse numbers in *Drosophila* looming responsive descending neurons

**DOI:** 10.1101/2024.04.24.591016

**Authors:** Anthony Moreno-Sanchez, Alexander N. Vasserman, HyoJong Jang, Bryce W. Hina, Catherine R. von Reyn, Jessica Ausborn

## Abstract

Synapses are often precisely organized on dendritic arbors, yet the role of synaptic topography in dendritic integration remains poorly understood. Utilizing electron microscopy (EM) connectomics we investigate synaptic topography in *Drosophila melanogaster* looming circuits, focusing on retinotopically tuned visual projection neurons (VPNs) that synapse onto descending neurons (DNs). Synapses of a given VPN type project to non-overlapping regions on DN dendrites. Within these spatially constrained clusters, synapses are not retinotopically organized, but instead adopt near random distributions. To investigate how this organization strategy impacts DN integration, we developed multicompartment models of DNs fitted to experimental data and using precise EM morphologies and synapse locations. We find that DN dendrite morphologies normalize EPSP amplitudes of individual synaptic inputs and that near random distributions of synapses ensure linear encoding of synapse numbers from individual VPNs. These findings illuminate how synaptic topography influences dendritic integration and suggest that linear encoding of synapse numbers may be a default strategy established through connectivity and passive neuron properties, upon which active properties and plasticity can then tune as needed.

## INTRODUCTION

Dendrites are intricate tree-like structures that receive the majority of synaptic inputs. However, we still know very little about the organization of synaptic inputs on dendritic trees and how this arrangement influences specific computations. A better understanding of how synaptic inputs are organized is crucial to understanding dendritic processing and will provide insight into how neural function becomes disrupted in neurodevelopmental diseases and neurodegenerative disorders with altered synapse organization (Fiala et al., 2002; Johnston et al., 2016; Luebke et al., 2010; Wang and Balice-Gordon, 2022).

At the level of individual synapses, synapse location can affect computation through the passive attenuation of dendritic potentials, where distal synapses have less of an impact at the spike initiation zone (SIZ) than proximal synapses (Rall, 1959; Spruston et al., 2016, 1994). On the other hand, some neurons compensate for this passive attenuation by tuning synaptic and/or dendritic properties to equalize the impact of each individual synapse at the SIZ. This phenomenon has previously been termed dendritic or synaptic democracy (Häusser, 2001; Poirazi and Papoutsi, 2020; Rumsey and Abbott, 2006; Timofeeva et al., 2008).

At the level of synapse populations, the topographic organization of synapses or synapse clusters with similar functional tuning can lead to non-linear summation of synaptic inputs through passive mechanisms such as shunting, or through the activation of active conductances (Branco et al., 2010; Kirchner and Gjorgjieva, 2022; Poirazi and Papoutsi, 2020; Stuart and Spruston, 2015; Takahashi, 2019).

While some progress has been made in characterizing and understanding these phenomena through experimental and computational studies, the underlying mechanisms and functional implications often remain elusive due to a lack of detailed dendritic morphology information and precise knowledge of synapse location and functional tuning.

To begin to address these limitations, researchers have recently turned to the model organism *Drosophila melanogaster*. Publicly available electron microscopy (EM) datasets of the *Drosophila* brain provide detailed information about neuron morphologies, synapse locations, and the presynaptic neurons to which these synapses belong (Scheffer et al., 2020; Zheng et al., 2018). Two recent studies have utilized these EM datasets, in combination with detailed biophysical modeling, to examine the impact of individual synapse location on dendritic trees in *Drosophila* neurons. Both studies showed that dendritic postsynaptic potentials in two different neuron types of *Drosophila* are normalized by compensatory voltage attenuation by the time they reach the SIZ, following the concept of synaptic democracy (Hafez et al., 2023; Liu et al., 2022). These computational studies predict important functional implications for the neurons they studied and demonstrate the importance of investigating dendritic integration at this level of detail. However, they did not consider input from multiple cell-types with different functional tuning and retinotopic organization or how synapses from differently tuned presynaptic neurons and populations are arranged on the dendritic tree and how this organization might influence computation.

Evidence for strategic synapse organization exists across species and many examples can be found in visual systems. At the local scale of a few synapses, inputs from particular visual features can be clustered according to their functional tuning (Kirchner and Gjorgjieva, 2022; Takahashi, 2019). Neighboring synapses on ferret L2/3 dendrites or macaque V1 dendrites receive inputs with similar orientation preferences (Scholl et al., 2017; Wilson et al., 2016). Synapses in mouse V1 are instead organized based on their retinotopic receptive fields (Iacaruso et al., 2017; Jia et al., 2010). Examples of synapse organization into dendritic maps also exist. In the *Xenopus* tadpole optic tectum, mouse visual cortex, and the locust giant movement detector (LGMD) neuron, different regions of individual dendrites are tuned to stimuli in different locations of the retinotopic input map (Bollmann and Engert, 2009; Iacaruso et al., 2017; Zhu and Gabbiani, 2016). Why organization strategies may differ, and the functional role of these organizations remains to be thoroughly evaluated.

We here used the detailed information provided by the full adult fly brain (FAFB) EM dataset (Zheng et al., 2018) and EM-data derived multicompartment models fitted to experimental data to investigate how retinotopically tuned presynaptic information is organized across dendrites and how this organization impacts integration in postsynaptic partners. In *Drosophila*, a behaviorally relevant role for retinotopic processing has begun to be elucidated in looming detection circuits that process and encode looming stimuli, the 2-D projections of objects approaching on a direct collision course (Dombrovski et al., 2023; Keleş and Frye, 2017; Morimoto et al., 2020; Wu et al., 2016). Detecting the location of an approaching threat is vital for the survival of any animal, making it a highly salient feature of looming circuits, and therefore a relevant model system to study how retinotopic information is processed. Whether synaptic topography plays a role in organizing retinotopic information flow in this system is, however, unknown.

Looming detection circuits involve distinct sensory processing layers, from photoreceptors in the compound eyes to the lobula complex. Photoreceptors capture luminance changes, and subsequent layers—including the lamina, medulla, lobula, and lobula plate—process and integrate visual information while maintaining the retinotopic information imposed by the spatial arrangement of the photoreceptors (Borst, 2014; Borst and Helmstaedter, 2015; Joly et al., 2016; Sanes and Zipursky, 2010; Wu et al., 2016). In the final processing layers of the lobula and lobula plate, populations of visual projection neurons (VPNs) encode specific features or feature combinations, such as the size or angular velocity of looming stimuli (Ache et al., 2019; Cowley et al., 2023; von Reyn et al., 2017, 2014). VPN dendrites receive retinotopic information in the lobula and lobula plate and project axons to the central brain, where they synapse onto descending neurons (DNs). DNs integrate both the feature (what) as well as the retinotopic (where) information that is encoded by VPNs to then select and drive appropriate motor responses, such as directional escape jumps (Card and Dickinson, 2008a, 2008b; Dombrovski et al., 2023; Namiki et al., 2018; von Reyn et al., 2014) or flight avoidance maneuvers (Kim et al., 2023; Namiki et al., 2018; Ros et al., 2024; Schnell et al., 2017). Retinotopy of VPN to DN synapses has been suggested to be encoded in the form of synapse number (synaptic weight) gradients that appear across DN dendrites (Dombrovski et al., 2023), rendering a DN more responsive to visual features located in a particular region of the fly’s visual field. It remains to be determined how synapse location and topography also play a role in how DNs process information from VPNs.

At the level of VPN populations, the clustering of synapses of a given VPN type on the dendritic tree could introduce nonlinearities to the processing of retinotopic VPN inputs (Poirazi and Papoutsi, 2020; Spruston et al., 2016). This has been suggested for other model systems where synapses with similar characteristics are clustered along the dendritic tree (Kirchner and Gjorgjieva, 2022; Takahashi, 2019; Ujfalussy and Makara, 2020). Clustering of synaptic inputs has indeed been suggested in the looming circuits of *Drosophila*, as synapses originating from different VPN types terminate in distinct regions known as glomeruli (Mu et al., 2012; Otsuna and Ito, 2006; Panser et al., 2016; Wu et al., 2016) and therefore also project to non-overlapping areas on the DN dendrites. However, the functional implications of this possible clustering have not been investigated.

At the level of individual VPNs, it is also unknown if retinotopic information encoded by neurons of a VPN type is translated into a clustering of synapses from the same neuron or an overall retinotopic organization of VPN synapses along the dendritic tree. This could considerably shape the functional implications of synaptic gradient maps. For example, a recent study of the LGMD neuron in locusts has demonstrated that a retinotopic mapping of excitatory inputs can significantly impact the ability of the neuron to discern fine stimulus differences (Dewell et al., 2022). Computational models investigating synaptic organization and dendritic integration have also often suggested that synaptic organization influences the computations a neuron performs (Gulledge et al., 2005; Poirazi and Papoutsi, 2020; Spruston, 2008; Tran-Van-Minh et al., 2015).

Using the FAFB EM dataset and full morphology biophysical models of DNs and their VPN inputs, we investigated the topography of VPN to DN synapses within the looming detection circuits of *Drosophila* and how this organization, or lack thereof, impacts the integration of synaptic inputs.

Our work suggests that DN morphology and passive properties establish a synaptic democracy across their dendrites that extends to all VPN populations. Further, our analyses and models suggest that VPNs do not organize their synapses in a retinotopic map but effectively place synapses to retain linear encoding of synapse numbers when multiple synapses are coactivated. We therefore provide important insights into how retinotopic synaptic gradient maps could be preserved despite the challenges of synaptic integration in individual dendritic processes. Lastly, our models provide a platform for investigating active integration mechanisms in the future.

## RESULTS

### VPN synapse clusters are distributed across DN sub-branches

Looming responsive DNs in *Drosophila* integrate retinotopic visual information from multiple VPNs. To uncover general principles for how VPN information is organized on DN dendrites, we selected five representative DNs demonstrated or predicted to be looming responsive for our analyses: DNp01, DNp02, DNp03, DNp04, and DNp06 (Dombrovski et al., 2023; Jang et al., 2023; Namiki et al., 2018; Peek, 2018; von Reyn et al., 2017, 2017). A connectivity analysis of the FAFB EM connectome (Zheng et al., 2018), shows that this population of DNs integrates looming information from six main VPN populations (**Figure 1B**) that have dendrites either only in the lobula (LC4, LC6, and LC22) or both in the lobula and lobula plate (LPLC1, LPLC2, and LPLC4) of the ipsilateral optic lobe. VPNs of the same population project their axons out of the optic lobe to distinct, non-overlapping central brain regions, called glomeruli, except for LC22 and LPLC4, which project to a shared glomerulus (**Figure 1**, Mu et al., 2012; Otsuna and Ito, 2006; Panser et al., 2016; Wu et al., 2016). Neurons of each VPN population are morphologically distinct (**Figure 1D**) and synapse onto multiple looming responsive DNs (**Figure 1B**).

**Figure 1.**
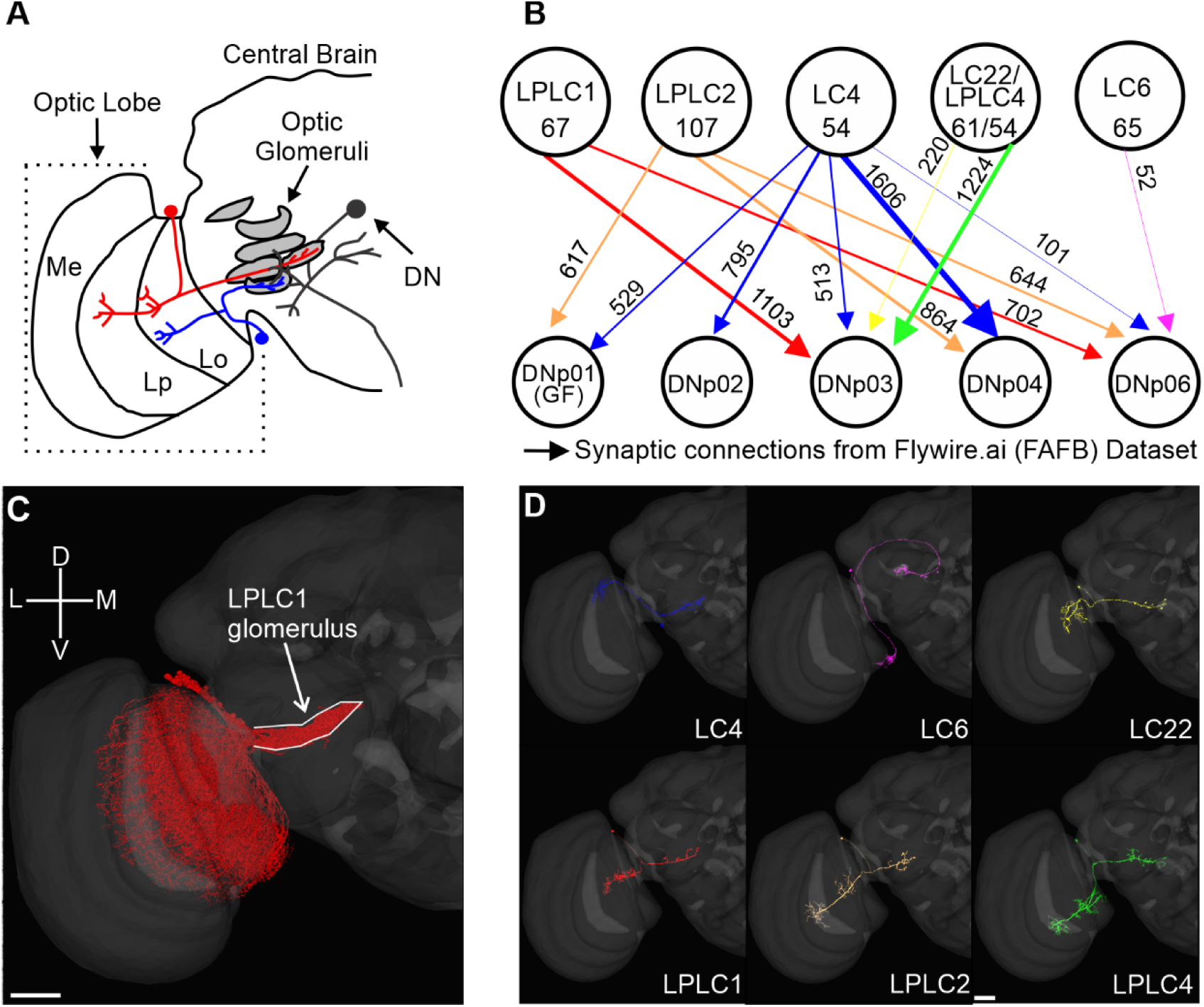
Populations of looming responsive VPNs synapse onto single ipsilateral DNs. (**A**) Schematic of the *Drosophila* optic lobe and central brain illustrating two VPNs from different populations that project to their respective glomeruli in the central brain where they synapse onto a DN. (**B**) Looming sensitive VPN connectivity onto five DNs based on the FAFB EM connectome. The numbers of neurons within a VPN population are given within the circles, and the numbers of synaptic connections made by the whole population onto each DN are given next to the connections. (**C**) EM reconstructed LPLC1 population from flywire.ai (Dorkenwald et al., 2022). (**D**) Single cell reconstructions of representative VPNs (LC4, LC6, LC22, LPLC1, LPLC2, and LPLC4). Scale bars 10 µm. Me, Medulla; Lp, Lobula plate; Lo, Lobula.

To investigate VPN-DN synaptic organization we first analyzed the broad topographic organization of different VPN population inputs on the DNs. Since individual VPN populations project their axons to dedicated non-overlapping optic glomeruli (Mu et al., 2012; Otsuna and Ito, 2006; Panser et al., 2016; Wu et al., 2016), synapses from the same population also projected onto non-overlapping regions of the dendrites of their DN synaptic partners (**Figure 2B-F, Left**). However, a dendrogram analysis showed that VPN synapse clusters were not localized on dedicated sub-branches of DN dendrites. Rather, synapses from different VPN populations generally projected to shared DN dendrite branches (**Figure 2B-F, Right**).

**Figure 2.**
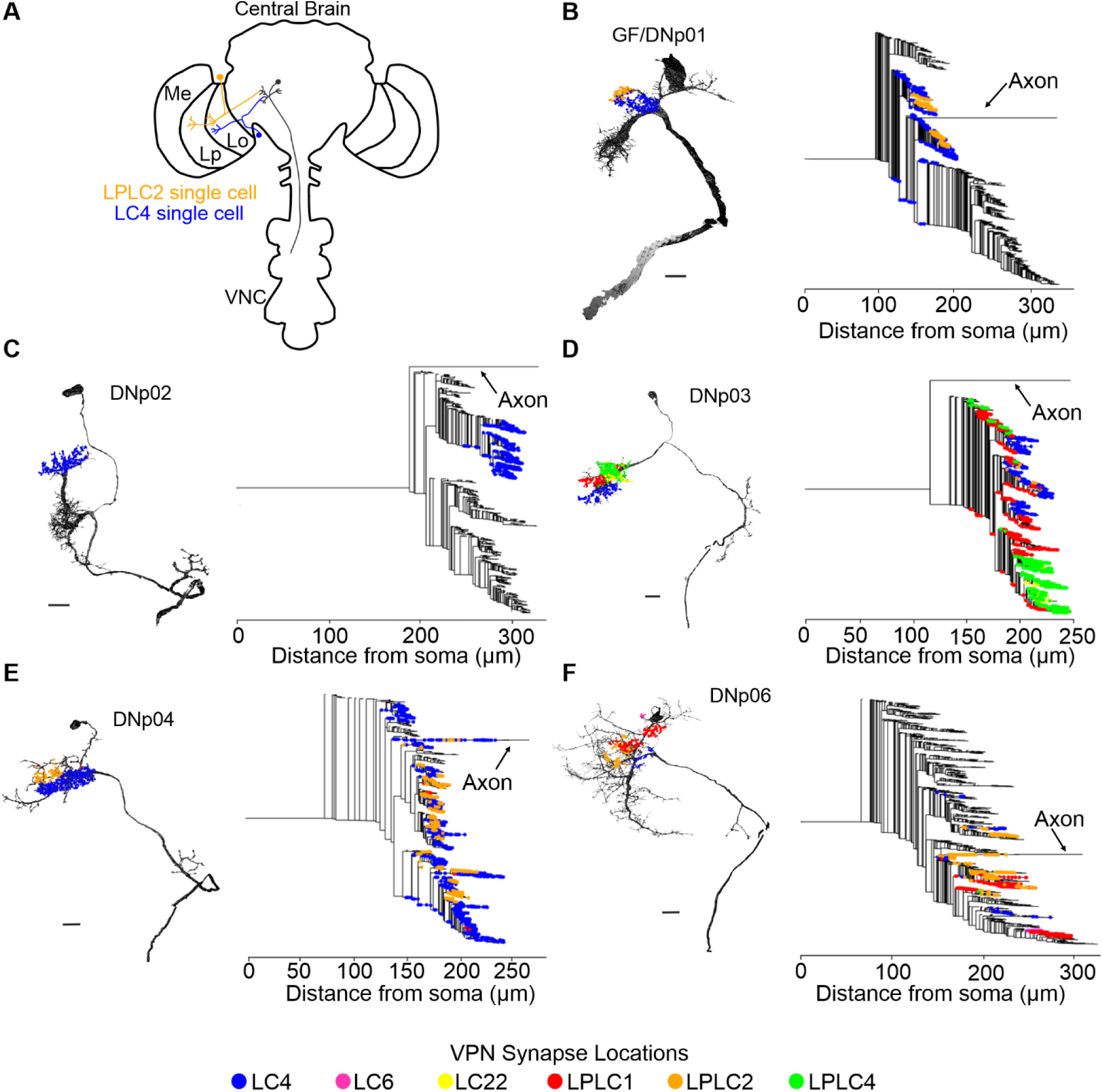
VPN inputs spatially cluster on DN partners but span multiple sub-branches. (**A**) Schematic of example VPNs from two populations (LC4 and LPLC2) synapsing onto a downstream partner (DNp01). (**B-F**) Left: Synapse locations of VPN populations across the dendritic arbors of the 3D EM reconstructions of five looming responsive DNs (DNp01, DNp02, DNp03, DNp04, and DNp06). VPN populations project their synapses to non-overlapping regions of the DN dendrites. Right: Dendrograms of the same five DNs with VPN synapse locations. VPN populations occupy overlapping sub-branches. Scale bars: 15 µm.

LPLC2 and LC4 synapse clusters, for example, are segregated along the distal-proximal axis of both DNp01 lateral dendrites (**Figure 2B, Left**) (Ache et al., 2019; McFarland et al., 2024). From the DNp01 dendrogram, we found synapses from the LC4 and LPLC2 populations were split evenly among the two distinct visual dendrite sub-branches (**Figure 2B, Right**). Similarly, for the other DNs, while VPN synaptic inputs were spatially segregated when visualized on the EM reconstructions, the dendrograms showed that sub-branches were shared across VPNs, and not dedicated to a single VPN type (**Figure 2D-F**). DNp02 was the exception in that it only received synapses from LC4, which synapsed onto one main dendrite (**Figure 2C**).

In summary, while synapses from different VPN populations project to non-overlapping regions, our analysis showed that they are not confined to separate dendritic sub-branches. These data suggest distinct VPN populations are therefore not separated into discrete electrotonic compartments.

### VPN synapses are not retinotopically organized

While each VPN cell type encodes a particular visual feature or combinations of visual features (Ache et al., 2019; Cowley et al., 2023; Klapoetke et al., 2022; von Reyn et al., 2017), they also provide the location of the feature(s) through the activation of individual neurons within each population (Dombrovski et al., 2023; Morimoto et al., 2020). The dendrites of individual VPNs project to different regions of the lobula and lobula plate where they receive retinotopic information from upstream partners (Morimoto et al., 2020; Wu et al., 2016). This dendritic organization can be leveraged to anatomically estimate the retinotopic receptive field of each neuron using EM data reconstructions of their dendrite branching patterns in the lobula (Dombrovski et al., 2023; Morimoto et al., 2020).

Previous work has demonstrated that the retinotopic organization of VPNs, and consequently the locations of visual stimuli, are conveyed to DN partners through synaptic gradients, wherein the number of synapses from VPN inputs to their postsynaptic DN partners increases along a particular axis of the fly’s visual field (Dombrovski et al., 2023). However, some but not all VPN populations also exhibit retinotopy in how their axons terminate within a respective glomerulus (Dombrovski et al., 2023; Morimoto et al., 2020). We, therefore, investigated whether retinotopic information is also preserved downstream through the organization of VPN synapses on their DN partners, which could further shape or amplify existing synaptic gradients.

Using VPN morphologies reconstructed from flywire.ai (Dorkenwald et al., 2022), we generated anatomical estimations of the receptive fields for the looming responsive populations LC4, LC6, LC22, LPLC1, LPLC2, and LPLC4 following a previously established methodology (Dombrovski et al., 2023; Morimoto et al., 2020). For each neuron within a given VPN population, we generated a polygon around their dendritic arbors. These polygons were then projected onto the curvature of the lobula to estimate the area of the visual field they occupy using known correlations between lobula and visual space (Dombrovski et al., 2023; Morimoto et al., 2020). We then calculated the centroid of each polygon as an estimate for the center of the neuron’s receptive field.

The resulting anatomical receptive fields are shown in **Figure 3**. Our estimations also corroborated two previously published estimated receptive fields that were generated using independent reconstructions of the LC4 (Dombrovski et al., 2023) and LC6 populations (Morimoto et al., 2020) from the FAFB dataset (Zheng et al., 2018). Although the VPN populations varied considerably in their number of neurons, ranging from 44 (LC22) to 108 (LPLC2) neurons per hemisphere, the collective dendrites of each population tiled the full visual field (**Figure 3**).

**Figure 3.**
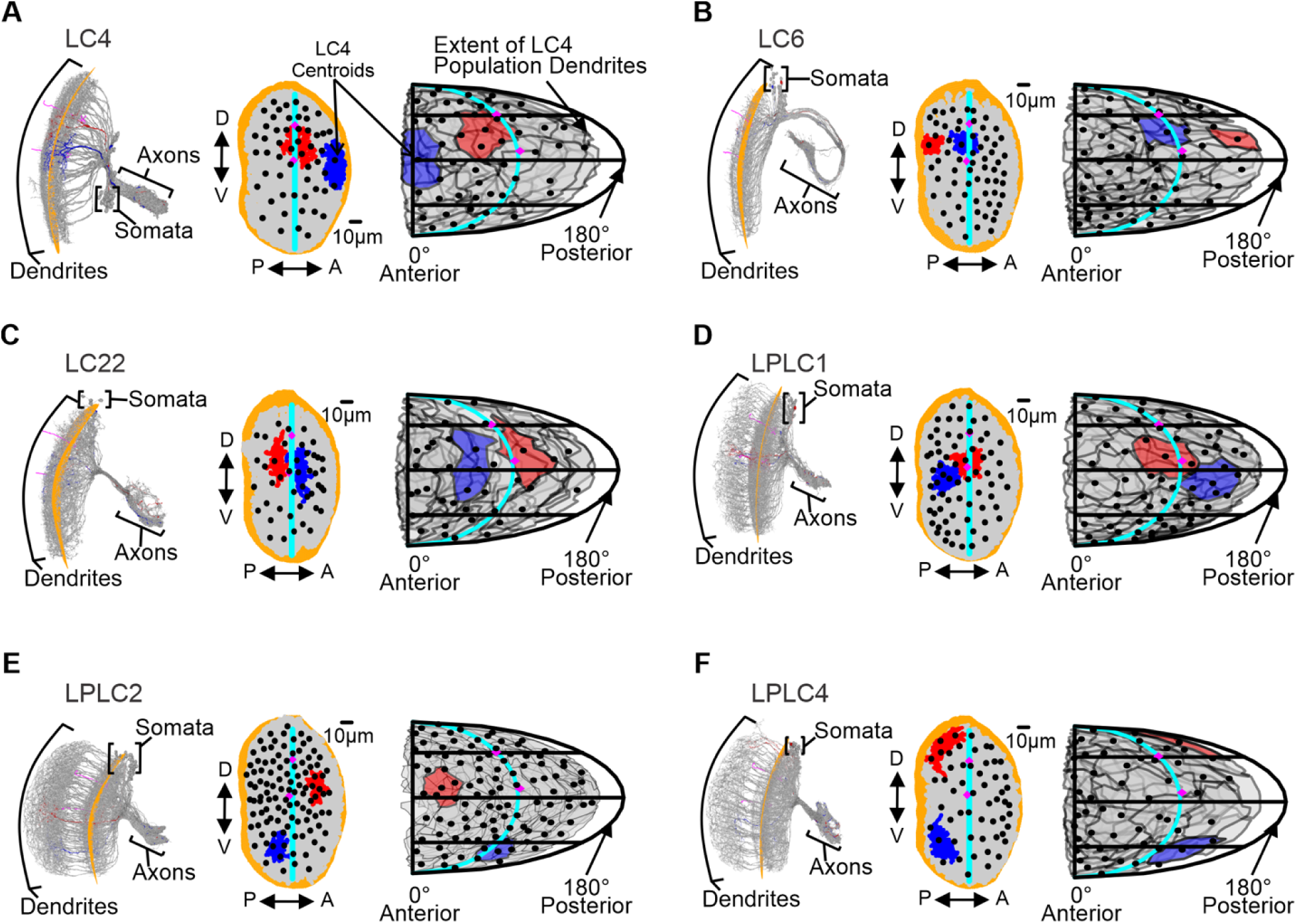
Anatomical receptive field estimations for looming responsive VPN populations tile the whole visual space. (**A**) Left: EM reconstructions of the LC4 population (gray) with a quadratic plane (orange) fitted to all LC4 dendrites to estimate the lobula curvature. Middle: 2D lobula projections of LC4 dendrites (gray) and centroids (black). Right: 2D estimation of dendritic receptive fields of the LC4 population projected onto the fly’s visual field. Translucent gray polygons represent the area of the visual field sampled by the dendrites of an individual neuron. (**B-F**) Same as in (A), but for LC6, LC22, LPLC1, LPLC2, and LPLC4. All Panels: Two example neurons are highlighted in red and blue. Two previously reconstructed Tm5 cells and their centroids are used as landmarks for the center of the eye and a more dorsal position of the central meridian (magenta). The central meridian is denoted by the cyan line. An analysis of how VPN centroids are organized within the lobula space is shown in **Figure 3—figure supplement 1**.

From the VPN centroids, we found that all assessed VPN populations presented slight biases in their distribution across one or both axes of the visual field. These biases were identified by assessing whether the centroids of a population were overrepresented in one hemisphere when dividing the lobula along either the anterior-posterior or dorso-ventral axis. LC4 centroids were more abundant in the dorsal hemisphere of the lobula. LC6, LC22, and LPLC4 centroids were denser in the anterior half of the lobula. LPLC1 centroids showed a bias toward the posterior hemisphere of the lobula, and LPLC2 centroids exhibited a bias toward the dorsal and posterior hemispheres (**Figure 3—figure supplement 1**).

Using the estimated receptive field centroids, we then investigated if and how different VPN populations preserved retinotopy while projecting their synapses onto postsynaptic partner DNs. We illustrate this on two specific VPN-DN pairs: LC4-DNp01 and LPLC4-DNp03 in **Figure 4**. These pairings were chosen because these VPN populations have previously been reported to exhibit axonal retinotopy and would therefore be more likely to also organize their synapses retinotopically (Dombrovski et al., 2023). Within these pairings, we visualized two groups of synapses belonging to VPNs located at the extremes of the dorso-ventral and anterior-posterior axes. If retinotopy was preserved, synapses from neurons at opposite extremes along an axis would project to distinctly different locations. Utilizing EM reconstructions of the DNs and their presynaptic VPN synapse locations, we found no apparent synapse retinotopy from the two extreme anterior-posterior and dorso-ventral VPN subsets in the LC4-DNp01 and LPLC4-DNp03 sample pairs (**Figure 4 A,B**).

**Figure 4.**
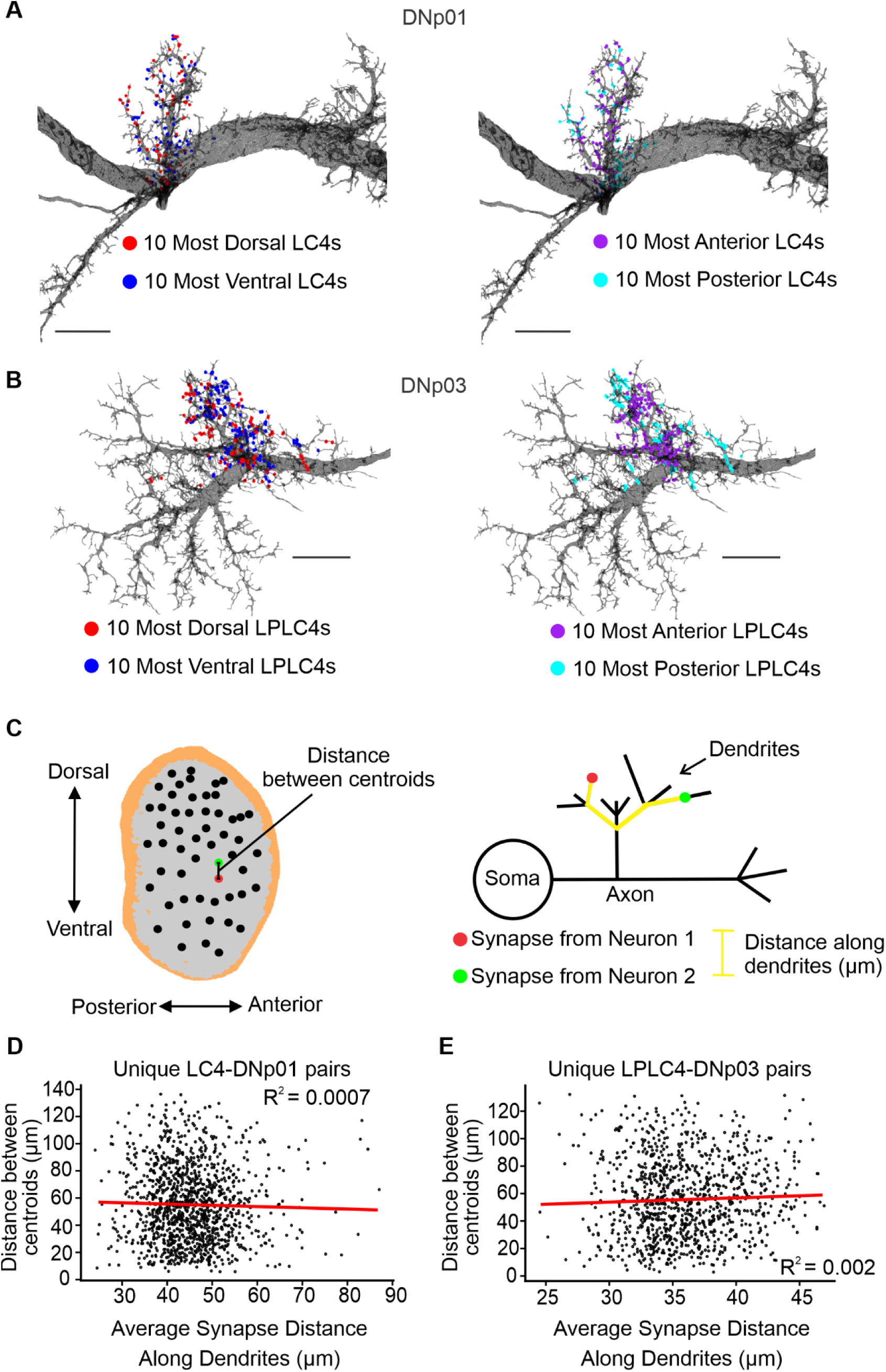
VPN synapses are not retinotopically organized. (**A,B**) Representative examples illustrating no retinotopic organization of two VPN subsets selected from the extreme positions on each lobula axis. (**A**) Left: Synapse locations of the 10 most dorsal and ventral LC4 neurons on the EM surface reconstruction of DNp01. Right: Synapse locations of the 10 most anterior and posterior LC4s. (**B**) Same as (A) but for LPLC4-DNp03. Scale bars: 15 µm. (**C**) Left: Schematic illustrating the calculation of Euclidean distance in the lobula space for a selected pair of neurons. Right: Schematic illustrating the calculation of the distance along the dendrites (µm) for a pair of synapses. For each unique pair of neurons, the average synapse distance is calculated as the average of the distances between all possible unique synapse pairs between the two neurons. (**D,E**) Representative plots show a lack of retinotopic synapse organization. A positive linear trendline would indicate retinotopically organized synapses. (**D**) The distance between the centroids of all unique pairs of LC4 presynaptic to DNp01 is not correlated with the average synapse distance for that given pair of LC4 neurons. Linear regression in red. (**E**) Same as (D) but for LPLC4-DNp03. Analyses as illustrated in (C) for all other VPN-DN pairs are shown in **Figure 4—figure supplement 1**.

To further investigate this relationship, we performed a linear regression analysis of the distance between unique VPN centroid pairs and the average distance between the synapses of the same two neurons. We, again, found no correlation for the LC4-DNp01 and LPLC4-DNp03 sample pairs (**Figure 4 A,B**) or any of the other VPN-DN pairings (**Figure 4—figure supplement 1**). If retinotopy was preserved at the synapse level, synapses from neurons with similar receptive fields (closer to each other in the lobula space) would also be located closer on average. However, no such correlation was observed, suggesting no retinotopic synapse organization for any of the investigated VPN-DN pairs.

Taken together, our analyses show that all investigated VPNs tile the complete lobula (and therefore also visual) space but show slight biases in their distribution along one or both of the main axes of the lobula space. In addition, none of the six investigated VPN populations organize their synapses on DN dendrites in a retinotopic manner.

### VPNs project synapses to neighboring locations

While VPN synaptic inputs onto DN partners are not retinotopically organized, a more local organization of synapse clusters with the same retinotopic tuning could exist at the level of a few synapses. Local (over tens of microns) clustering of synapses with similar functional tuning that lack a more global organization, is emerging as a common organizing principle across brain regions and species (Kirchner and Gjorgjieva, 2022). This could have implications for processing since experimental evidence in several model systems suggests that simultaneous activation of local synapse clusters could lead to dendritic nonlinearities through passive or active mechanisms (Kirchner and Gjorgjieva, 2022; Takahashi, 2019).

Utilizing the fact that synapses from the same neuron inherently have the same tuning, we were able to use the FAFB EM dataset to test for local functional clustering of synapses by analyzing if two nearest neighbor (NN) synapses are more likely to belong to the same neuron than not.

To this end, we generated ten shuffled synapse datasets for each VPN-DN pair by shuffling the presynaptic synapse IDs, resulting in datasets where the locations of synapses were preserved, but their presynaptic identities were scrambled (**Figure 5A**). For all unique pairs of NN synapses, we then determined the percentage of NN pairs where both synapses belonged to the same presynaptic neuron. A comparison of the percentages for the original dataset and an average of the ten shuffled datasets showed that NN pairs belonged to the same neuron at significantly higher rates (∼50 %) than in the shuffled datasets (1–9 - 7.9 %) (Mann–Whitney U test, ****p < 0.0001) (**Figure 5B**).

**Figure 5.**
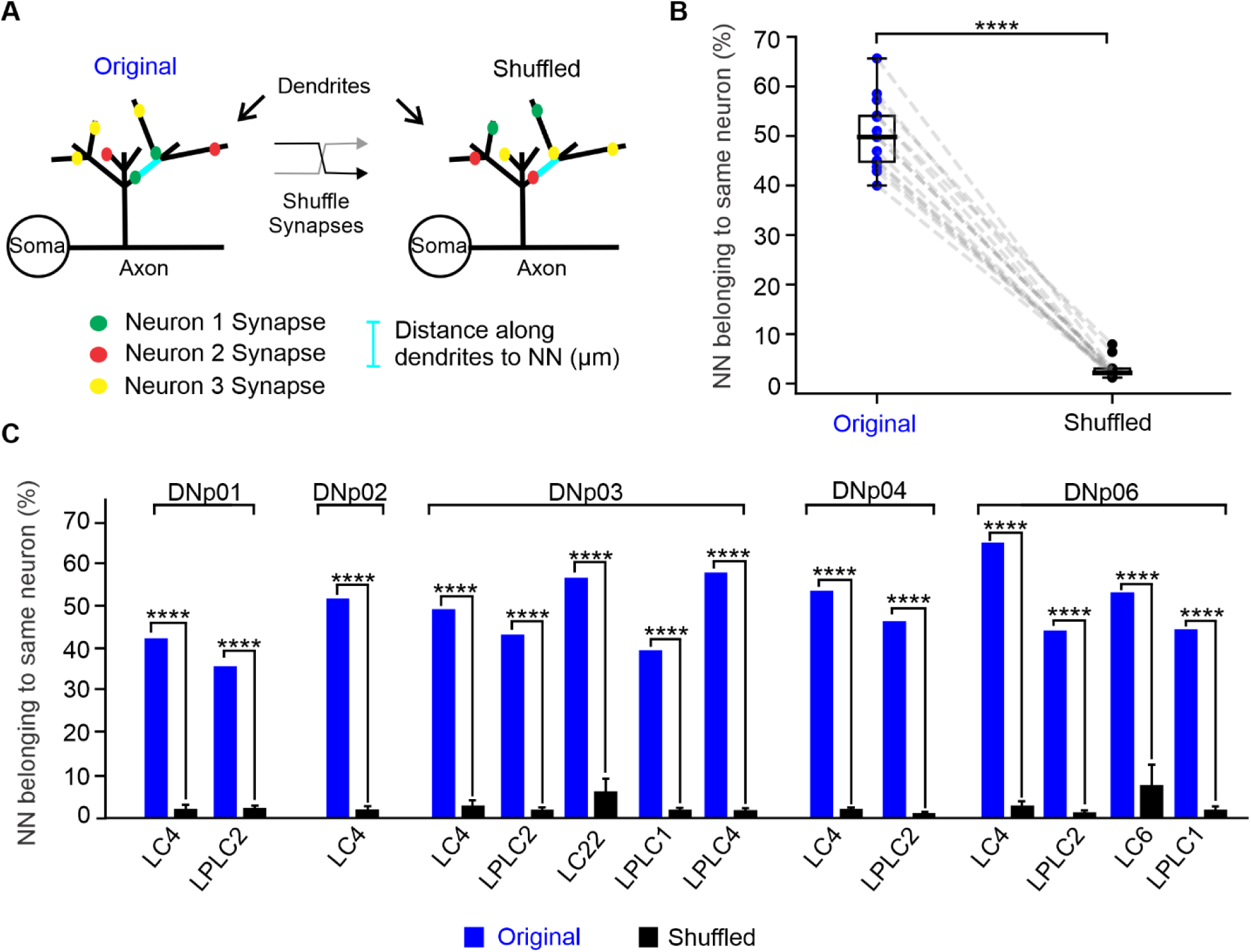
VPNs project their synapses to neighboring locations. (**A**) Schematic representation of classifying nearest neighboring (NN) synapse pairs along the DN dendrites in the original dataset and after shuffling synapse identities. (**B**) Percentage of unique NN synapse pairs belonging to the same neuron for all VPN-DN pairs for the original and shuffled datasets, Mann–Whitney U test, ****p < 0.0001 (**C**) Percentage of unique NN synapse pairs belonging to the same neuron for each VPN-DN pair. 1-sample T-test ****p < 0.0001.

Therefore, although VPN synapses do not show retinotopic organization on DN dendrites, synapses from the same VPN—and thus also synapses with the same functional tuning—tend to cluster together at the local scale of a few synapses. We note here that these small local clusters do not indicate all synapses from a single neuron form tight cohesive clusters onto the same dendritic region. To better understand how this organization impacts overall integration, and how it compares to tightly clustered versus random synapse distributions, we turned to computational modeling.

### Passive multicompartment models from EM data

To investigate the impact of VPN-DN synaptic organization on synaptic integration, we developed multicompartment computational models of DNs with accurately mapped VPN synaptic inputs. Using the flywire.ai (Dorkenwald et al., 2022) environment for the FAFB EM dataset (Zheng et al., 2018), we retrieved surface mesh reconstructions of the five looming responsive DNs (DNp01, DNp02, DNp03, DNp04, DNp06) as well as coordinate information describing the locations of their presynaptic looming responsive VPN input synapses (LC4, LC6, LC22, LPLC1, LPLC2, LPLC4) along the dendritic arbors of these DNs.

We developed a simple pipeline (**Figure 6**) to convert the EM surface mesh reconstructions into skeleton models while preserving biological radius information and then mapped the synapse locations onto the resulting skeletons. After skeletonizing and proofreading the EM meshes, the skeletons were exported as a series of connected nodes into NEURON via Python, and multicompartment DN models were generated using the coordinates, radius information, and parent-child hierarchy defining the individual model sections. Synapses were then mapped onto the models. The resulting multicompartment DN models with biologically accurate VPN synapse location placement were then used to simulate synapse activations.

**Figure 6.**
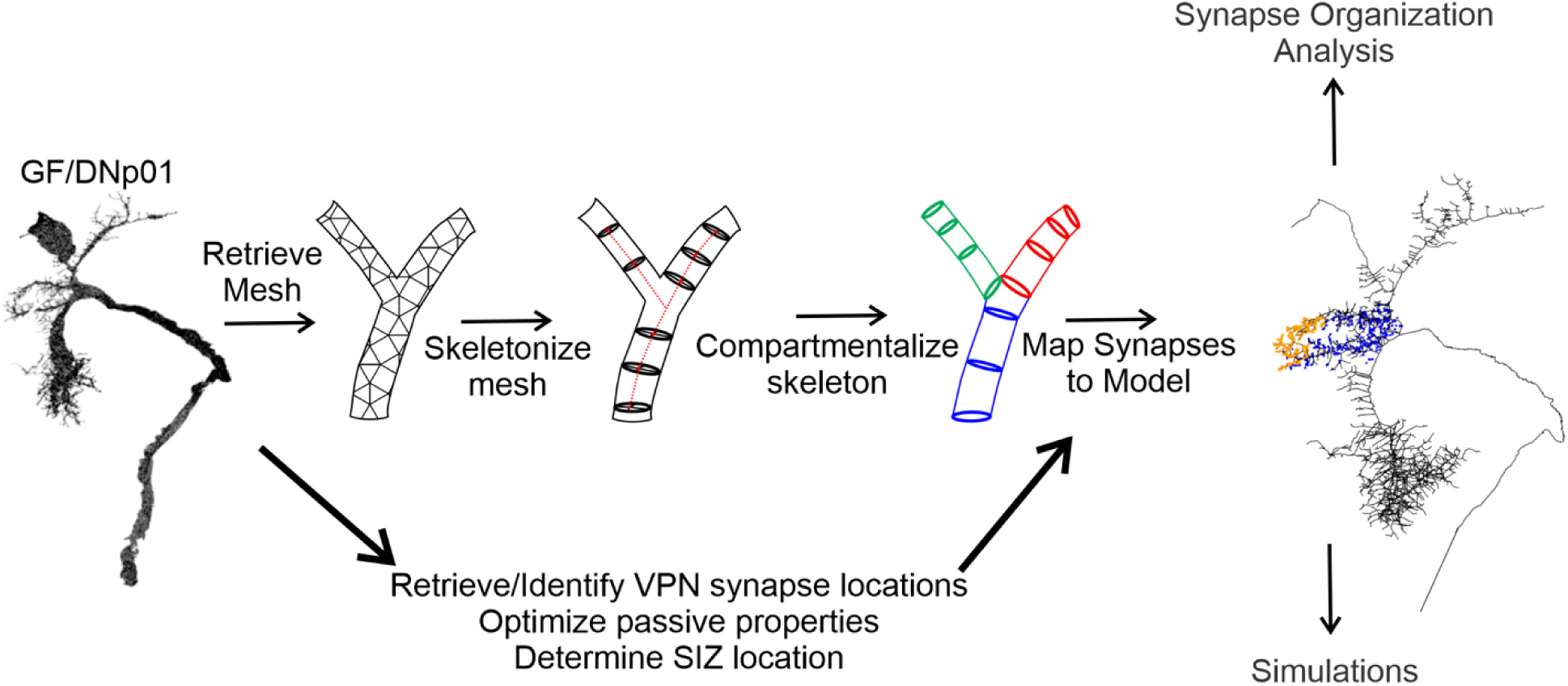
Overview of the modeling pipeline.

Passive properties for two representative DN models (DNp01 and DNp03) were tuned to fit experimental data from whole-cell, current clamp recordings from the soma of each DN (**Figure 7**). DNp01 was selected for its unique morphology with large diameter processes. DNp03 was selected for its more representative morphology and because it receives a majority of the VPN inputs, including the DNp01 inputs LC4 and LPLC2 (**Figure 1**). Small current injections were used to ensure that no voltage-gated ion channels were activated, and that the membrane responded passively across trials. The resulting model parameters [capacitance (*C_m_*), leak conductance (*g_leak_*), axial resistivity (*R_a_*), and reversal potential (*E_rev_*)] were found to be within physiological ranges (Borst and Haag, 1996; Gouwens and Wilson, 2009) and corresponded to other published models of *Drosophila* neurons (Gouwens and Wilson, 2009; Günay et al., 2015; Hafez et al., 2023; Liu et al., 2022).

**Figure 7.**
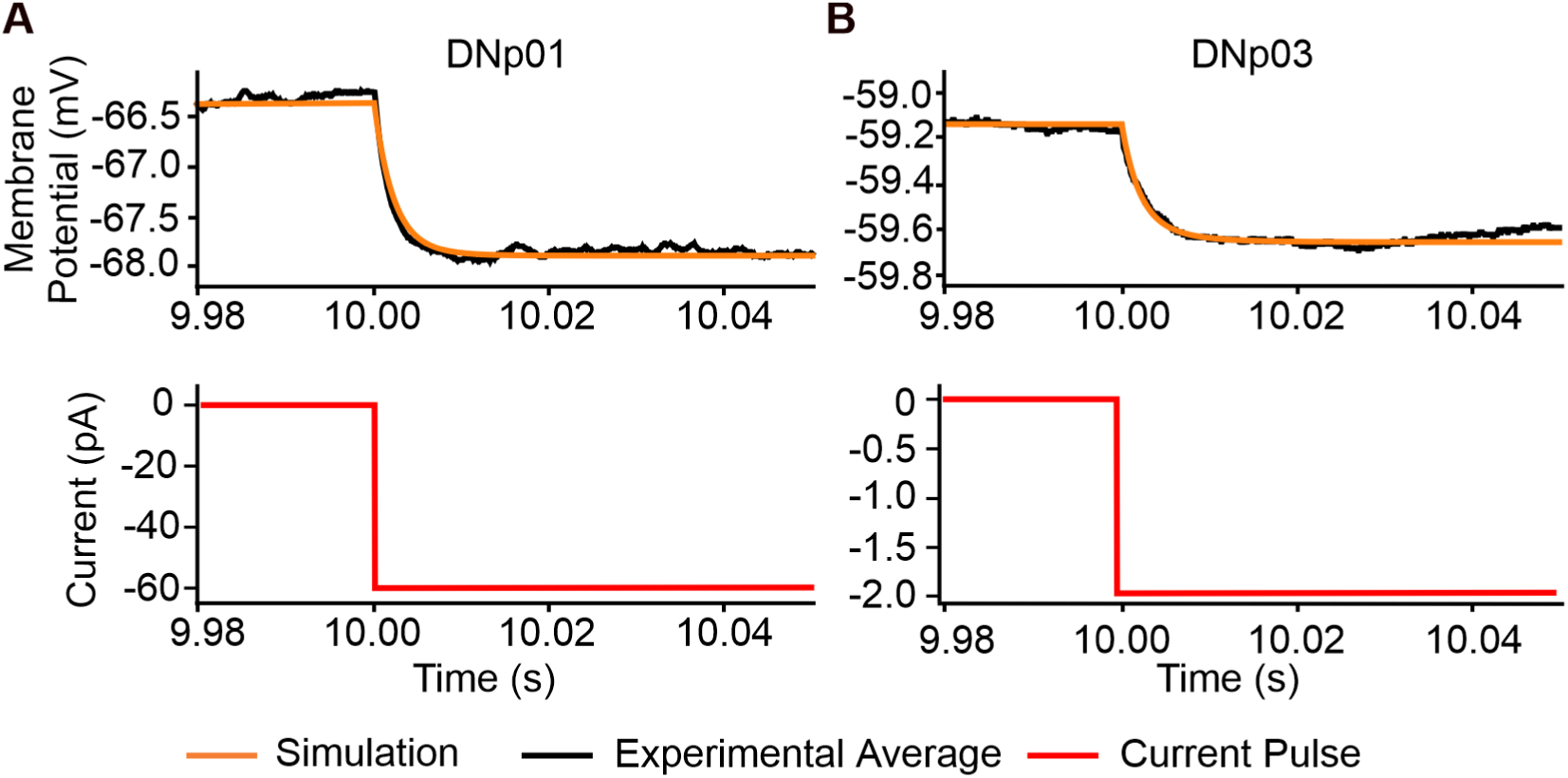
Fitting passive properties of the models. (**A**) Simulated current injection into the soma of a multicompartment model of DNp01, fitted to averaged experimental current injections (−60 pA) from three animals, 20 sweeps per animal. (**B**) Same as (A) but for DNp03 (current injection: −2 pA).

All 5 investigated DNs possess the capacity for action potential generation (Dombrovski et al., 2023; Jang et al., 2023; Namiki et al., 2018; Peek, 2018; von Reyn et al., 2017, 2014). These DNs therefore are predicted to feature a spike initiation zone (SIZ) characterized by a high concentration of voltage-gated sodium channels, whose principal alpha subunit is encoded by the *para* gene (Loughney et al., 1989). Since the SIZ is the final point of synaptic integration, we determined the SIZ location in each of the DNs. We utilized the split-GAL4 system to express RFP within a DN of interest (Namiki et al., 2018) and *para*-GFSTF (Ravenscroft et al., 2020) to label the endogenous expression of *para* with GFP throughout the whole brain. This allowed us to use confocal microscopy to identify regions of each DN that highly express voltage-gated sodium channels (**Figure 8**).

**Figure 8.**
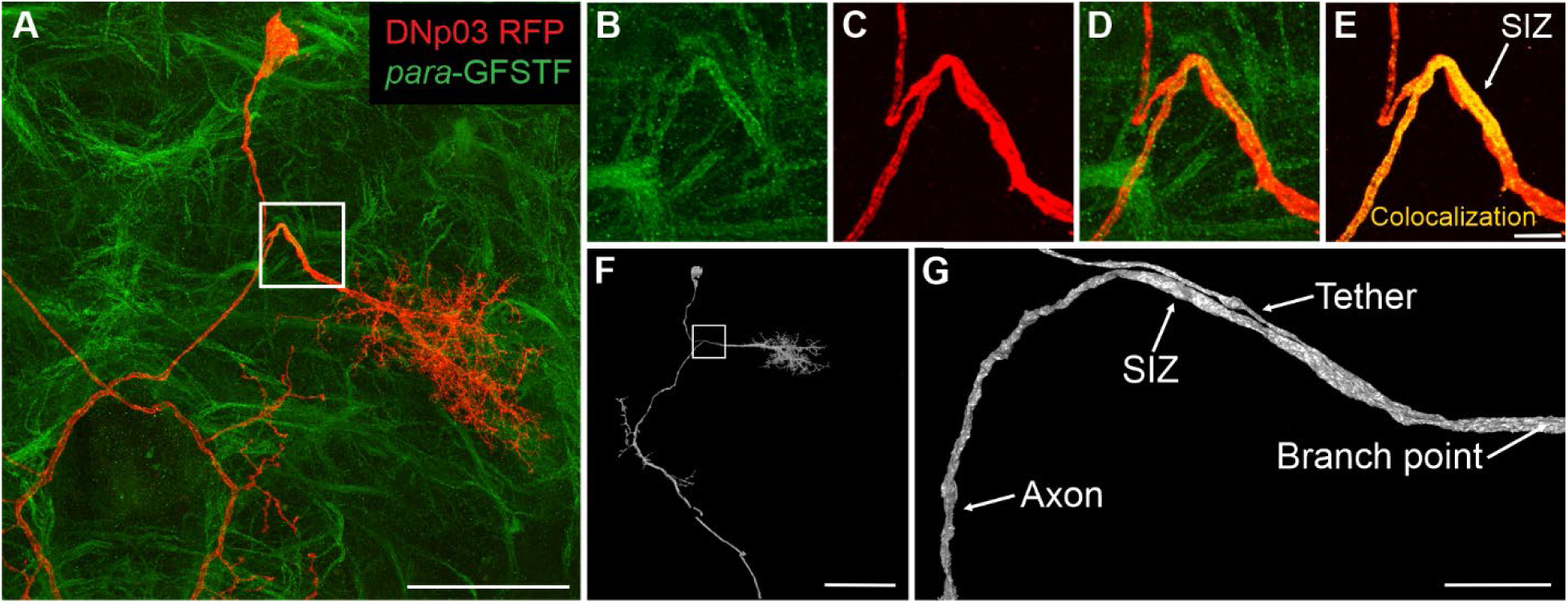
The SIZ is located downstream of the tether in DNp03. (**A**) Maximum intensity projection of RFP labeled DNp03 and GFP labeled para (genotype: *para-GFSTF, DNp03-split-GAL4, UAS-RFP*). Scale bar: 50 µm. (**B-E**) Zoomed-in view of (A) showing, (**B**) the individual para, and (**C**) DNp03 channels, (**D**) their overlay, and (**E**) para colocalization on DNp03. Scale bar: 5 µm. (**F**) FAFB EM mesh reconstruction of DNp03 (ID: 720575940627645514). Scale bar: 50 µm. (**G**) Zoomed-in view of the DNp03 mesh reconstruction from (F) showing the tether branchpoint region. Scale bar: 7.5µm. SIZ locations for the other DNs are shown in **Figure 8—figure supplement 1-3**.

For all evaluated DNs, we found the SIZ is consistently located downstream of the morphological branch point where the dedicated soma tether connects to the axon (**Figure 8E**, **Figure 8—figure supplement 1-3**). For DNp03 we found that the tether runs parallel to the axon before connecting, resulting in this junction occurring closer to the dendrites (**Figure 8G**).

### DNs passively normalize VPN synaptic potentials

Our analyses of synapse organization showed that VPN synapses generally cover a large range of DN dendrite locations (**Figure 2**) and would therefore be subject to varying levels of passive attenuation until they reach the SIZ, where they are integrated (Häusser, 2001). This is also a factor to consider when comparing synapses from different VPN populations, which project to non-overlapping dendritic regions (**Figure 2**) and therefore might also be subjected to different levels of passive attenuation. This organization could affect the relative integration of excitatory postsynaptic potentials (EPSPs) from different VPN types or from individual neurons by introducing a proximal to distal bias.

While passive attenuation and differences in distal to proximal synapse effectiveness have been well described for some neurons (Poirazi and Papoutsi, 2020; Spruston et al., 2016), previous experimental and computational studies have shown that, in some neurons, the amplitude and time course of postsynaptic potentials are independent of synapse location (Hafez et al., 2023; Häusser, 2001; Liu et al., 2022; Otopalik et al., 2019, 2017; Timofeeva et al., 2008). This location independence has been termed synaptic democracy and both active as well as purely passive mechanisms have been proposed (Häusser, 2001; Jaffe and Carnevale, 1999; London and Segev, 2001; Otopalik et al., 2019; Rumsey and Abbott, 2006; Timofeeva et al., 2008; Williams and Stuart, 2003). Using our detailed multicompartment models of DNs we investigated how individual synapse location affects EPSP amplitude at the SIZ, as the site of final integration of information.

Using VPN inputs to DNp01 and DNp03 as representative examples, we simulated the activation of individual VPN synapses and recorded the resulting EPSP both at the synapse location and at the SIZ (**Figure 9**). All synapses were implemented with identical conductances and kinetics. Our simulations showed that although the initial EPSP amplitude at the synapse location varied widely from synapse to synapse (**Figure 9**; DNp01: 0.2-5 mV; DNp03: 0.2-1.7 mV), EPSP amplitudes at the SIZ were normalized to a narrow range (**Figure 9**; DNp01: 0.045-0.061 mV; DNp03: 0.16-0.19 mV). While EPSP amplitudes at the SIZ for DNp03 were larger than for DNp01, both were within the range of previous models of *Drosophila* neurons (Hafez et al., 2023; Liu et al., 2022) or large neurons from other model systems (Häusser, 2001; Magee and Cook, 2000; Winters et al., 2017). Within the narrow amplitude range at the SIZ, we observed a slight bias in amplitudes correlated with the synapse distance to the SIZ. Synapses proximal to the SIZ had slightly larger amplitudes compared to more distal synapses (**Figure 9**). This distance dependent bias has also been noted in other *Drosophila* neurons that exhibit synaptic democracy (Hafez et al., 2023; Liu et al., 2022).

**Figure 9.**
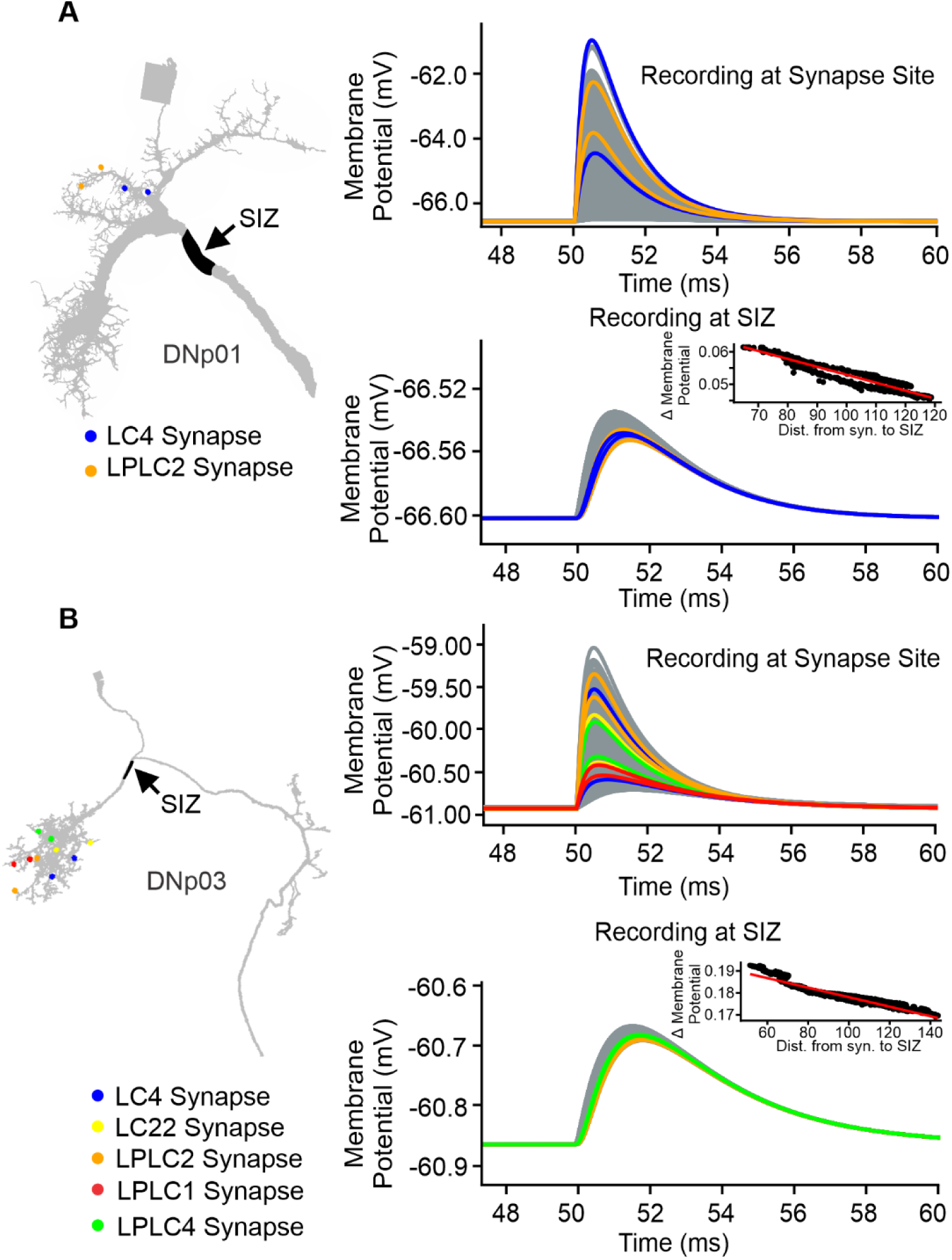
VPN synaptic input is passively normalized at the SIZ. (**A**) Left: Schematic of the DNp01 model with the SIZ location in black and randomly selected example synapses from each VPN population color coded. Right: Membrane voltage traces from activations of individual VPN synapses. Example synapse EPSPs are color coded accordingly. All remaining VPN synapse EPSPs are shown in gray. Top: Membrane voltage at the synapse location. Bottom: Membrane voltage of the same simulations at the SIZ. Inset: EPSP amplitude at the SIZ of individual VPN synapses over their distance to the SIZ. (**B**) Same as (A), but for DNp03 and its VPN inputs.

Our results suggest DN morphologies and their passive properties are tuned to normalize the effectiveness of individual VPN synapses independent of their location on the dendrite, similar to previous reports of synaptic democracy in *Drosophila* and other animal models (Hafez et al., 2023; Häusser, 2001; Jaffe and Carnevale, 1999; Liu et al., 2022; Otopalik et al., 2019, 2017; Timofeeva et al., 2008; Williams and Stuart, 2003).

### Passive DN models linearly encode VPN synapse numbers

Our multicompartment models predict that DNs normalize VPN synapse EPSPs, establishing a synaptic democracy as previously suggested for other *Drosophila* neuron types (Hafez et al., 2023; Liu et al., 2022). We next aimed to better understand how this effect translated to the passive integration of multiple synaptic inputs. Although synapses appear normalized when activated individually, nonlinear integration may occur when multiple synapses are activated simultaneously based on their topography (Kirchner and Gjorgjieva, 2022; Tran-Van-Minh et al., 2015). To understand how the synapse topography of individual VPN neurons may affect their integration at the SIZ, we activated individual VPNs using their precise synapse locations.

For both DNp01 and DNp03, we simultaneously activated all synapses of individual VPN neurons and found that the composite EPSPs at the SIZ increased linearly with the number of synapses (**Figure 10**). This linear encoding of synapse numbers did not depend on VPN type as the activations resulted in similar depolarizations for all VPN types presynaptic to the same DN (**Figure 10**). These data suggest, when only passive properties are taken into consideration, the impact an individual VPN neuron has on a DN response is directly correlated to the number of synapses it makes with a DN.

**Figure 10.**
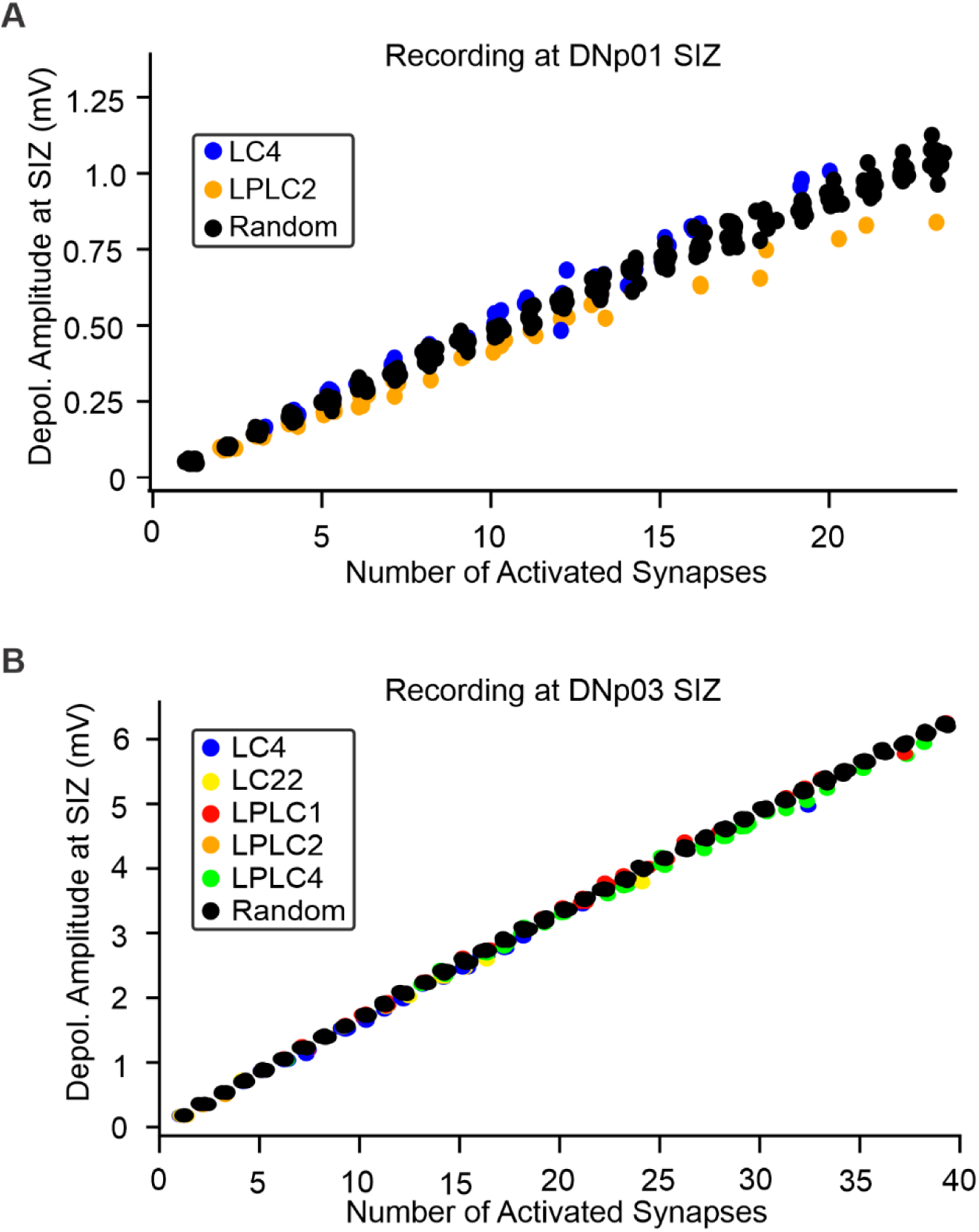
Linear encoding of VPN synapse numbers. (**A**) Depolarization amplitude at the SIZ of DNp01 in response to the activation of multiple synapses. Color-coded circles: Activation of all synapses of an individual VPN. Black circles: Activation of randomly selected synapse sets. (**B**) Same as (A) but for DNp03 and its VPN inputs.

We found nearest neighboring synapse pairs from the same neuron were more likely to belong to the same neuron than not (**Figure 5**). It is therefore possible that local clustering of synapses from an individual neuron causes a net shunting effect. To investigate potential shunting effects from local clustering, we compared the composite EPSP amplitudes at the SIZ when activating all synapses of an individual neuron with those of randomly selected synapse sets. These random synapse sets (which would be more spread out over the dendrite than synapses from the same neuron) resulted in the same depolarization at the SIZ as actual neuron activations (**Figure 10**), confirming that no, or very little shunting affects the integration of these synapses. Our results suggest that although individual neurons show a bias towards clustering their synapse locations, synapse distributions still retain enough spread to limit shunting.

We next investigated whether linear encoding of synapse numbers is retained when higher numbers of synapses are activated simultaneously. For each VPN population presynaptic to DNp01, we randomly selected and activated an increasing number of synapses until the whole population was activated.

The simulations showed that, while synapse numbers are encoded linearly for the simultaneous activation of synapses from individual neurons, activation of increasingly higher synapse numbers led to sublinear summation through shunting of the EPSPs (**Figure 11**). For DNp01 inputs, shunting was more pronounced for LPLC2 than LC4 at the same number of synapses (**Figure 11A**). For DNp03, the amplitude dependence and appearance of shunting were similar between populations (**Figure 11B**).

**Figure 11.**
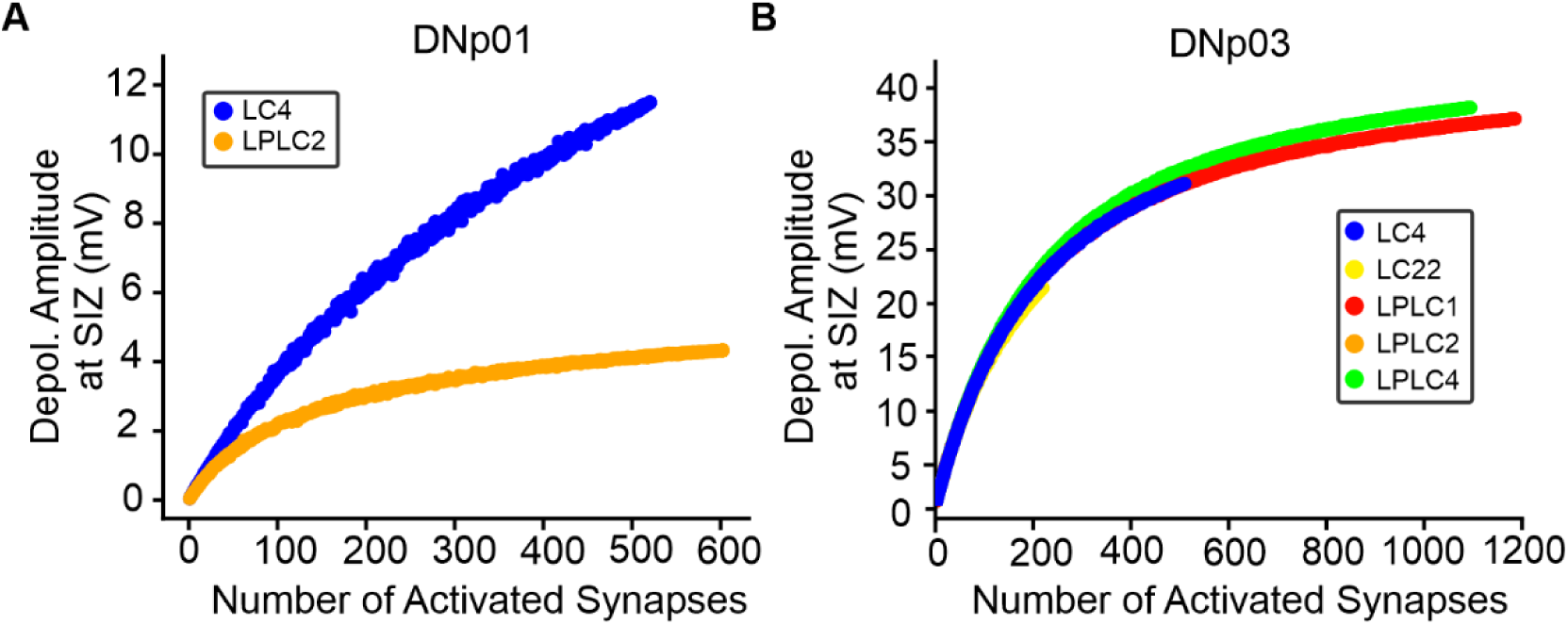
Shunting of VPN inputs with increasing synapse activation. (**A**) Peak depolarization at the SIZ of DNp01 in response to the activation of an increasing number of synapses from a given VPN population. (**B**) Same as (A) but for DNp03 and its VPN inputs.

Together this suggests that DN dendrites passively integrate their synaptic inputs sublinearly, and linear encoding of synapse numbers is only maintained for the activation of fewer than 100 synapses (corresponding to the activation of only a few neurons).

### Distribution of VPN synapses across DN dendrites prevents shunting

The observation that VPN synaptic inputs only operate in a linear range for the activation of synapses from a few neurons (**Figure 10**, **Figure 11**), led us to further investigate whether the actual distribution of VPN synapses is operating in an ideal range to reduce shunting effects. We first considered two extreme cases by simulating synapse distributions where all synapses are in close proximity (which should maximize shunting) and synapses randomly distributed across all VPN dendrites (which should reduce shunting).

We randomly selected *n* VPN synapses (random group), and a second distribution where we selected a random synapse and its n-1 nearest neighbors (close group) and compared their distributions to a single VPN (**Figure 12 A,B**). For each DN, we generated 50 synapse sets for the two groups (random and close) and simulated their simultaneous activation, recording the composite EPSPs at the SIZ (**Figure 12**).

**Figure 12.**
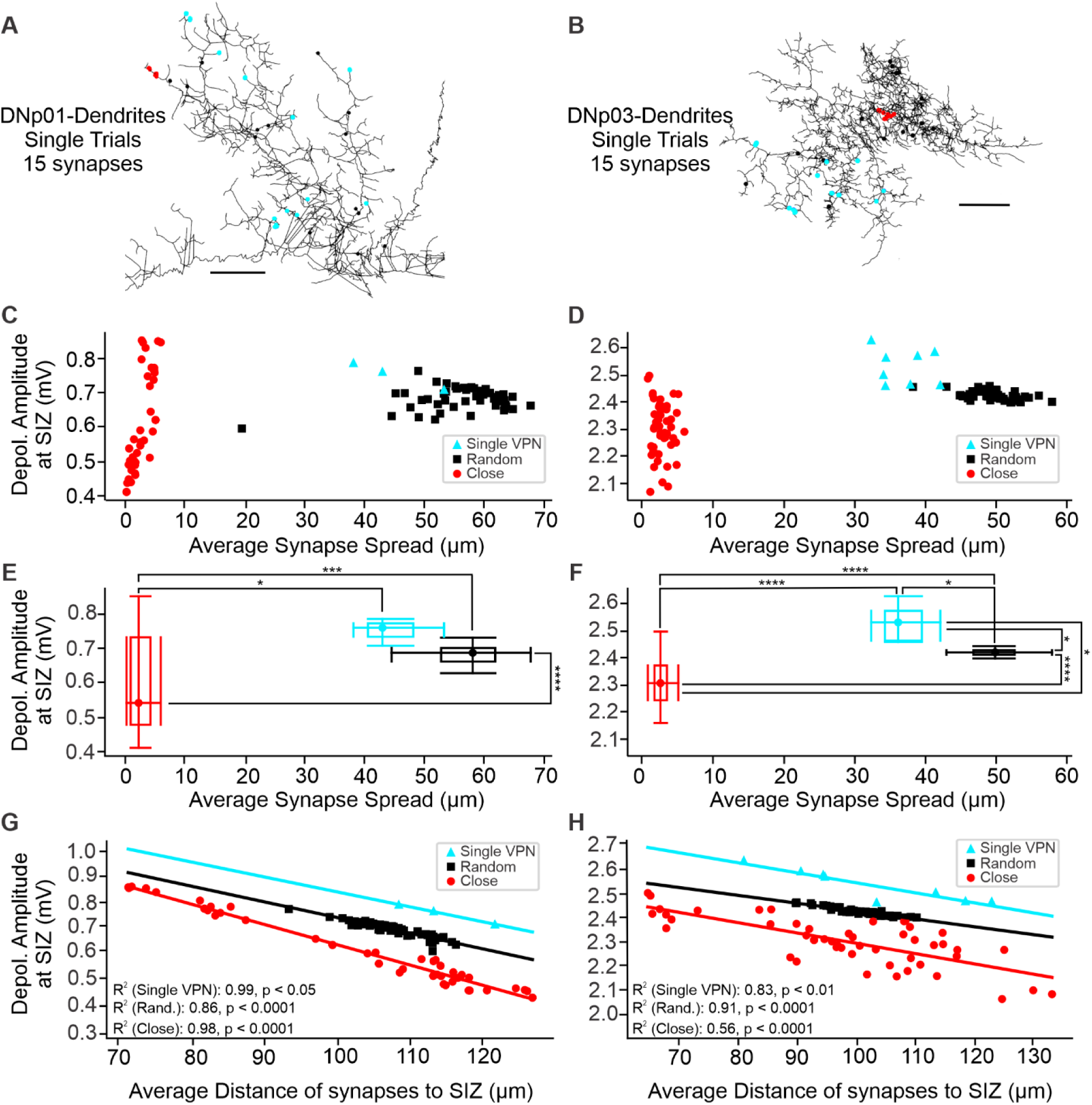
Individual VPNs distribute synapses across DN dendrites to avoid shunting. (**A**) Example synapse locations across the DNp01 skeleton model for single sets of 15 close and random synapses and for an example VPN with 15 synapses. (**B**) Same as (A) but for DNp03. Scale bars: 15 µm. (**C, D**) Composite EPSP amplitudes for the simultaneous activation of the three groups of synapse sets. Composite EPSP amplitudes cover a wider range for the Close group, than for the Random or Single VPN groups. (**E, F**) Population analysis of data from (C) and (D) shows VPNs distribute their synapses in a random fashion to maximize composite EPSP amplitudes (Kruskal-Wallis tests p < 0.05, followed by a Dunn’s post-hoc test with a Bonferroni adjustment of p-values *p <0.05, **p < 0.01, ***p <0.001, ****p < 0.0001)(**G, H**): Composite EPSP amplitude depends on the average synapse distance to the SIZ. The Close group covers a broader range of distances to the SIZ and therefore also a larger range of EPSP amplitudes than the other two groups. Analysis for a range of synapse numbers is shown in **Figure 12—figure supplement 1**.

We found, when activating the same number of synapses, that the close group showed a significantly wider range of composite EPSP amplitudes than the random group for both DNp01 (Levene’s Test: p < 0.0001) and DNp03 (Levene’s Test: p < 0.0001) (**Figure 12 C-F**). These data suggest a randomized synapse distribution is an ideal strategy to enable a linear relationship between the number of synapses activated and the composite EPSP amplitude at the SIZ. We found this trend was consistent across a range of synapse numbers (2-20 synapses) (**Figure 12—figure supplement 1**).

To better understand the observed differences between random and close synapse topography, we evaluated the composite EPSP amplitudes over the average distance of the synapses to the SIZ. We found randomizing synapse locations decreased the range of average distances to the SIZ when compared to the close synapse group. We also observed a distance-dependent covariation of the composite EPSP amplitudes that could explain the large EPSP variation in the close group (**Figure 12 G,H**)—the impact of activating a population of close synapses decreases with distance from the SIZ.

We then asked where activations of actual VPN synapse locations fall in relation to the close and random distributions. Interestingly, we found when activating actual VPN with the same number of synapses in our models (15 synapses for the examples shown in **Figure 12**), their responses are no different than random in terms of composite EPSP amplitude for DNp01, and even slightly larger in DNp03 (Kruskal-Wallis tests p < 0.05, followed by a Dunn’s post-hoc test with a Bonferroni adjustment of p-values, *p< 0.05).

Together, our data suggest that despite local clustering of synapses from the same neurons (**Figure 5**), VPNs distribute their synapses enough to avoid strong shunting effects to interfere with the linear encoding of synapse numbers.

## DISCUSSION

The organization of synaptic inputs on dendritic trees plays a critical role in neural computation, yet its functional implications remain poorly understood. In this study, we utilized detailed morphological data from the full adult fly brain (FAFB) electron microscopy (EM) dataset and biophysical computational models fitted to experimental data to investigate the organization of synaptic inputs from looming responsive visual projection neuron (VPN) populations onto their downstream descending neuron (DN) partners in *Drosophila melanogaster*. We find both DN morphology and the organization of VPN synapses on DN dendrites enable a linear encoding of synapse numbers. This property of DN passive integration may serve as a default neuronal blueprint upon which active properties and synaptic plasticity can build.

### Synapse topography of feature information

While VPN feature encoding populations spatially cluster their synapses in glomeruli (Keleş and Frye, 2017; Klapoetke et al., 2022; Wu et al., 2016), we find these clusters do not coincide with dedicated dendritic subbranches of the postsynaptic partners (**Figure 2**). This suggests that while synapses from different VPN populations likely don’t interact directly, distinct VPN populations are also not separated into continuous electrotonic compartments. This patchwork organization of smaller VPN clusters, that becomes apparent when considering the electrotonic structure revealed by the dendrograms, might serve to avoid strong nonlinearities that have been described for clusters of synapses with the same functional tuning. For example, in the layer 2/3 (L2/3) pyramidal neurons of the ferret primary visual cortex, the coactivation of synapses that are spatially clustered and have similar orientation tuning contributes to robust input–output nonlinearities in a neuron’s response to inputs from across its dendritic field, which is suggested to diversify orientation selectivity (Wilson et al., 2016). Similarly, in the L2/3 pyramidal cells of the mouse V1 primary visual cortex, synaptic inputs representing similar visual features and overlapping receptive fields are more likely to terminate on nearby dendritic branches, which has been suggested to facilitate nonlinear dendritic integration (Jia et al., 2010; Longordo et al., 2013).

While synapses from distinct VPNs are not organized in continuous clusters, they nonetheless are organized into smaller clusters on the different subbranches (**Figure 2**). This segregation may allow for distinct computations to be performed on each feature, potentially utilizing differential ion channel expression patterns to facilitate distinct nonlinear computations for specific features and shape dendritic integration during the activation of multiple input types (Hardie and Pearce, 2006; Häusser, 2001; Rumsey and Abbott, 2006; Williams and Stuart, 2003). The actual distribution of ion channels in these dendrites, however, is unknown and will be the subject of future studies.

### Synapse topography of retinotopic information and linear encoding of synapse numbers

We found no evidence of retinotopic organization of VPN synapses on DN dendrites (**Figure 4**) despite the clear retinotopic arrangement of VPN dendrites (**Figure 3**). We found neither a tendency for retinotopic organization along the two main axes (**Figure 4 A,B**), nor when we analyzed synaptic retinotopy by taking the distance along the dendrite into account (**Figure 4 D,E**). This was surprising since EM reconstructions of VPN morphologies showed that for some VPN populations, dendrite retinotopic organization is preserved in the organization of VPN axons as they enter their respective glomeruli (Dombrovski et al., 2023; Morimoto et al., 2020; Ribeiro et al., 2018; Wu et al., 2016). The functional relevance for the retinotopic organization of axons has yet to be established.

The lack of synaptic retinotopic maps, however, suggests a functional relevance for the distribution of synapses across the dendrites, since the formation and extension of axon collaterals into a wider net is resource intensive (Gallo, 2011). Therefore, supported by our models, we predict that the lack of retinotopy on DN dendrites serves a purpose and is part of a wiring design strategy to avoid shunting effects caused by the coactivation of neighboring synapses (**Figure 12**). By dispersing synapses, the individual potential of each synapse can be maximized and maintained when multiple synapses are coactivated. This allows a direct relationship between the number of synapses and the impact on the postsynaptic DN. Retinotopic gradients in synapse numbers have indeed been found in many VPN-DN pairings that induce directional behaviors in response to looming stimuli (Dombrovski et al., 2023). Allowing synaptic gradients to guide directional behaviors may necessitate the distribution of synapses across the dendrites, thereby avoiding dendritic nonlinearities that could confound the direct encoding of stimulus location.

Retinotopic synapse maps have been shown to exist on dendrites in different model systems (Bollmann and Engert, 2009; Iacaruso et al., 2017; Kirchner and Gjorgjieva, 2022; Zhu et al., 2018). It is important to note, however, that previous studies often analyzed retinotopic organization along an axis in physical space (similar to **Figure 4 A,B**) but not in dendrite space (measuring distances along the dendritic processes, **Figure 4 C-E, Figure 4—figure supplement 1**), since this type of analysis is only possible when information about both the exact synapse locations and the morphology of the dendrite is available. How these retinotopic maps translate to the electrotonic structure of the dendrites remains to be investigated.

Additionally, our models showed that encoding becomes sublinear for larger numbers of synapses (**Figure 11**). This suggests that DNs can effectively integrate a small number of synapses, but the integration becomes less efficient for larger synapse numbers. We currently have no solid estimates of how many neurons would be co-activated by a natural looming stimulus, but our anatomical receptive field analyses (**Figure 3**), along with recent Ca^2+^ imaging receptive field mappings (Klapoetke et al., 2022), suggest only a handful of neurons would be active in response to a looming stimulus, keeping the encoding in a near-linear range.

Our simulations also have implications for optogenetic experiments. Since shunting already occurs at relatively low synapse numbers (**Figure 11**), optogenetic stimulation would likely not be suited to investigate quantitative changes in synapse density at the level of VPN populations. It may instead be more appropriately used to differentiate the presence or absence of direct synaptic connections or estimate synaptic density when only limited cell numbers are activated.

We found the activation of all synapses from a single VPN neuron in both DNp01 and DNp03 resulted in higher composite EPSP amplitudes at the SIZ, reducing shunting effects in comparison to the close group, and in DNp03 also summating more efficiently than the random group **(Figure 12)**. This suggests that VPN synapses may be distributed on DN dendrites to take advantage of variations in dendrite diameters and the extensive branching structures of DNs to maximize their spatial summation while avoiding local shunting (Mocanu et al., 2000; Tran-Van-Minh et al., 2015).

While we observed no global retinotopic maps for VPN synapses, we found that synapses from the same VPN tended to cluster together at the local scale (**Figure 5**), suggesting a functional clustering of synapses with similar tuning. Local synapse clusters have been described as near-ubiquitous across brain areas and species (Kirchner and Gjorgjieva, 2022) and our analyses also predict their existence in *Drosophila* DNs, a synaptic site of sensorimotor integration (Namiki et al., 2018; Wu et al., 2016). Clusters of synapses with similar functional tuning play a crucial role in dendritic integration by influencing neuronal input-output transformations and enhancing computational capabilities (Kirchner and Gjorgjieva, 2022; Leighton et al., 2023; Poirazi and Papoutsi, 2020; Takahashi, 2019; Ujfalussy and Makara, 2020). They have also been suggested to be essential for high-fidelity signal processing and memory allocation (Kastellakis et al., 2015; Kastellakis and Poirazi, 2019). Local synapse clusters can facilitate the induction of active properties such as dendritic spikes and plateau potentials, which can enable advanced computational capabilities within dendrites of single neurons (Kastellakis and Poirazi, 2019; Poirazi and Papoutsi, 2020; Ujfalussy and Makara, 2020). The formation of these clusters likely occurs through local synaptic interactions (Kastellakis et al., 2015; Kirchner and Gjorgjieva, 2022; Larkum and Nevian, 2008; Leighton et al., 2023).

However, at least for our passive models, these clusters did not translate into strong nonlinearities due to shunting (**Figure 10**) and our analysis even shows that VPNs distribute their synapses in a near-random manner when considering all synapses of an individual neuron (**Figure 12 E,F**). We have yet to explore possible functional implications of these small clusters in *Drosophila* neurons with realistic active properties.

### Synaptic democracy in DNs

Our computational models demonstrate that DN morphologies passively normalize the effectiveness of individual VPN synapses, resulting in a synaptic democracy across their dendrites (**Figure 9**). This is consistent with previous studies that demonstrated synaptic democracy in *Drosophila* and other animal models (Hafez et al., 2023; Häusser, 2001; Liu et al., 2022; Otopalik et al., 2019, 2017; Williams and Stuart, 2003).

Some studies have suggested that amplifying distal synaptic conductances or shaping synaptic potentials through dendritic ion channels could be how synaptic democracy is established (Häusser, 2001; Häusser and Mel, 2003; Jaffe and Carnevale, 1999; Rumsey and Abbott, 2006, 2006; Williams and Stuart, 2003). On the other hand, the idea that synaptic democracy can emerge from passive electrotonic structure alone has gained accumulating evidence from demonstrations of synaptic democracy in passive models with detailed morphologies and experimentally supported SIZ locations and passive properties (Hafez et al., 2023; Liu et al., 2022; Otopalik et al., 2019, 2017; Timofeeva et al., 2008).

In our models, we assumed uniform conductances for all synapses, consistent with recent models of *Drosophila* neurons (Hafez et al., 2023; Liu et al., 2022; Tobin et al., 2017). Treating each synapse equally in models of *Drosophila* neurons is supported by EM data analyses of *Drosophila* synaptic contact site surface areas, which were found to be comparable across different brain regions and synapse types, and can be seen as an anatomical proxy for synaptic conductance (Barnes et al., 2022; Liu et al., 2022; Scheffer et al., 2020; Tobin et al., 2017).

DNs integrate retinotopic feature information from multiple VPN types that project to non-overlapping regions on the dendrite. In this type of arrangement, synaptic democracy could serve to remove the confounding factor of unequal synapse distance to the SIZ for the different VPN populations (and the features they encode). Further, organizing synapses in non-retinotopic maps together with democratizing synaptic amplitudes, could be a way to simplify the integration of retinotopic information that is encoded by the number of synapses each VPN makes with the postsynaptic DN (Dombrovski et al., 2023).

### Spike initiation zones in DNs

We experimentally determined the location of the spike initiation zone (SIZ) in the DNs, sometimes also called the axon initial segment. By using the split-GAL4 system to express RFP within the DN of interest (Namiki et al., 2018) and a transgenic *para*-GFSTF line (Ravenscroft et al., 2020) to label the endogenous expression of *para* with GFP, we showed that the SIZ is consistently located on the axon past the point where the soma tether connects to the axon (**Figure 8** and **Figure 8—figure supplement 1-3**). This was particularly striking for DNp03 where the soma tether runs parallel to the axon for a considerable distance (∼30 µm,) before connecting (**Figure 8G**). We therefore confirmed the existence of a SIZ for these spiking DNs and determined their location for use in our simulation studies.

Our results are also in qualitative agreement with previous work that identified the SIZ as a distal axonal segment in both larval and adult *Drosophila* neurons (Ravenscroft et al., 2020). Our experimental SIZ labeling also corroborates results from an earlier computational modeling study which estimated the putative SIZ location in *Drosophila* central neurons by simulating spike waveforms in computational models and determined that the SIZ would be located past the tether (Gouwens and Wilson, 2009).

An understanding of the location of spike initiation is crucial for the investigation of neuronal activity and for interpreting electrophysiological data accurately in invertebrates like *Drosophila* as the only structure accessible for patch-clamp electrophysiology is the soma, which is electrotonically distant from the SIZ and the dendritic regions (Gouwens and Wilson, 2009; Günay et al., 2015; Hafez et al., 2023; Liu et al., 2022). Furthermore, the precise anatomical location of the spike initiation zone significantly influences the input-output behavior of neurons, impacting their computational performance and information-processing capabilities (Günay et al., 2015; Verbist et al., 2020).

### Outlook

While our investigations of synaptic organization and integration are limited to the visual inputs of VPNs onto DNs, access to the EM connectome along with our pipeline will allow these investigations to be extended to other neural circuits. Our work also highlights the importance of synaptic organization, even in passive dendrites, to achieve efficient computations at the level of a single neuron. Our models now provide a foundation and computational framework to understanding how synaptic organization may interplay with a neuron’s intrinsic properties. Using these models together with experimental activation and manipulation of retinotopic processing in looming detecting circuits of *Drosophila* may provide important insights into more complex non-linear integration mechanisms in future studies.

## MATERIALS AND METHODS

### Experimental protocols

*Drosophila melanogaster* were raised on standard cornmeal fly food at 25°C and 60% humidity on a 12-h light:12-h dark cycle. For electrophysiology and immunohistochemistry experiments, female *Drosophila* that were between 2–5 days old post-eclosion were used. The following fly stocks were used for electrophysiology and immunohistochemistry experiments.

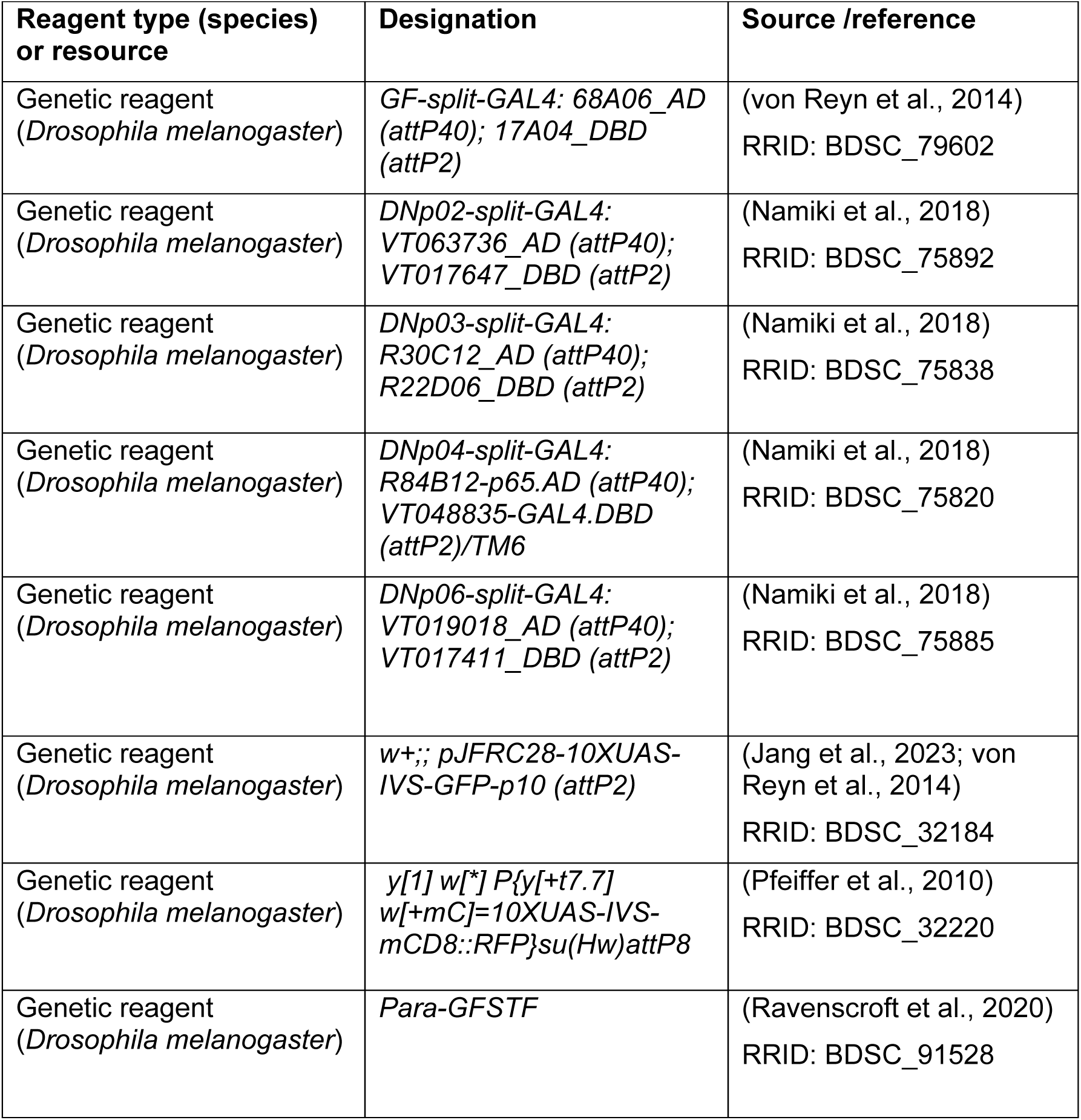

### Electrophysiological characterization of DNs

In vivo whole-cell, current-clamp electrophysiology was conducted on tethered flies as previously described (Jang et al., 2023; von Reyn et al., 2017). Flies were anesthetized by lowering the temperature to 4°C and their head and thorax were fixed to polyether-ether-ketone (PEEK) plates with UV glue (Loctite 3972). To access the DNs for electrophysiology, the cuticle and trachea were removed on the posterior side of the head, specifically in the area right above the DN soma, and the brain was perfused with standard extracellular saline (NaCl 103 mmol, KCl 3 mmol, TES 5 mmol, trehalose·2H_2_O 8 mmol, glucose 10 mmol, NaHCO_3_ 26 mmol, NaH_2_PO_4_ 1 mmol, CaCl_2_·2H_2_O 1.5 mmol and MgCl_2_·6H_2_O 4 mmol). (Gouwens and Wilson, 2009). Osmolarity was adjusted to 270–275 mOsm and the extracellular saline was bubbled with 95% O_2_/5% CO_2_ to maintain a pH of 7.3. All experiments were conducted at room temperature (20–22°C). The brain sheath was disrupted by applying collagenase (0.5% in extracellular saline) via an electrode and the region above the targeted soma was cleaned before recording. Patch-clamp electrodes (3-6 MΩ) were used to target GFP labeled DN somata and the electrodes were filled with intracellular saline (potassium aspartate 140 mmol, KCl 1 mmol, HEPES 10 mmol, EGTA 1 mmol, Na_3_GTP 0.5 mmol, MgATP 4 mmol, Alexafluor-568 5 µmol, 265 mOsm, pH 7.3). In vivo whole-cell recordings were performed in current-clamp mode using a MultiClamp 700B amplifier and digitized (NI-DAQ, National Instruments) at 20 kHz and low-pass filtered at 6 kHz. All data were obtained using the open-source software Wavesurfer (https://wavesurfer.janelia.org/) running in MATLAB (MathWorks). Traces were not adjusted for a 13 mV liquid junction potential (Gouwens and Wilson, 2009). Recordings were considered acceptable when the initial seal resistance was >2 GΩ before rupture and the resting membrane potential was less than −50 mV.

To determine passive membrane properties for each DN, 20 small hyperpolarizing pulses (< 2 mV) per animal for three animals were injected into each DN. Current amplitude, pulse duration, and time between pulses were adjusted for each DN to ensure they did not invoke any active currents and permitted full recovery to baseline. The resulting traces were then averaged across animals using MATLAB and used for passive property fitting in the models.

### Immunohistochemistry

All dissections were performed in cold Schneider’s insect media (S2, Sigma Aldrich, #S01416). Brains were then fixed overnight at 4°C in 1% paraformaldehyde (20% PFA, Electron Microscopy Sciences, #15713). Immunohistochemistry was performed as described previously (McFarland et al., 2024). Brains were then washed in phosphate buffered saline (Thermo Fisher Scientific Cat# 70011044) with 0.1% Triton X-100 (Sigma-Aldrich Cat# T8787) and incubated overnight at 4°C in with primary antibodies Rabbit-anti-dsRed (1:200, Takara Bio Cat# 632496, RRID:AB_10013483) and Chicken-anti-GFP (1:1000, Thermo Fisher Scientific Cat# A10262, RRID:AB_2534023). Brains were then washed and incubated overnight at 4°C with secondary antibodies Goat-anti-Chicken 488 (1:400, Thermo Fisher Scientific Cat# A-11039, RRID:AB_2534096) and Goat-anti-Rabbit 568 (1:400, Thermo Fisher Scientific Cat# A-11011 (also A11011), RRID:AB_143157). Brains were then washed and mounted onto coverslips coated with poly-L-lysine (Sigma Aldrich, #25988-63-0), dehydrated in increasing alcohol (EtOH) concentrations, cleared in Xylene (Fisher Scientific, #X5-500) as described previously (McFarland et al., 2024), and mounted in DPX mounting medium.

### Confocal microscopy

All images were acquired using a Leica Stellaris 5 confocal system. Images were taken with a 60x, 1.40 NA oil immersion objective. Imaging parameters were minimally adjusted between images to achieve an image that utilizes the full pixel intensity range without oversaturating pixels.

### Spike initiation zone estimation

To determine the location of the spike initiation zone (SIZ), or axon initial segment, for each DN, the default auto threshold in FIJI (RRID:SCR_002285) (Schindelin et al., 2012) was used to first generate a mask of the RFP-labeled neuron membrane. In MATLAB, the binarized membrane mask was then multiplied to the raw *para* channel pixel-wise for the entire stack of images. The resulting image was then generated representing the raw *para* intensity only where it colocalized with the neuronal membrane. The SIZ was indicated by a high density of colocalized *para* signal within the axon of the neuron of interest.

### Morphological reconstruction

#### EM mesh acquisition

Morphological reconstructions of *Drosophila* DNs were acquired from the ‘Full Adult Fly Brain’ FAFB dataset (Zheng et al., 2018) hosted in the Flywire database (Dorkenwald et al., 2022; https://edit.flywire.ai/, version 630; DNp01: “720575940622838154”, DNp02: “720575940619654053”, DNp03: “720575940627645514”, DNp04: “720575940604954289”, DNp06: “720575940622673860”). The FAFB dataset provides detailed neuron morphologies, with coordinates at 4 x 4 x 40 nm pixel resolution. For model implementation, coordinates were converted to microns to align with the default coordinate specification used by NEURON (Hines and Carnevale, 2001), a simulation environment designed for creating and simulating data-based model neurons with complex morphologies and their synaptic connections.

FAFB reconstructions of each DN include the full central brain morphology but do not contain the ventral nerve cord (VNC) portions of DN axons, as that anatomical region exists outside the bounds of the FAFB dataset. Axonal morphological data were gathered from the female adult *Drosophila* VNC electron microscopy dataset (FANC) (Phelps et al., 2021). The VNC dataset contains neuron morphologies with coordinates at 4.3 x 4.3 x 40 nm pixel resolution. Axonal reconstructions were obtained via the Virtual Fly Brain (https://fanc.catmaid.virtualflybrain.org, DNp01: “648518346477750423”, DNp02: “648518346523998725”, DNp03: “648518346517438244”, DNp04: 648518346490893859”, DNp06: “648518346479137937”). All meshes were obtained from the left side of FAFB (Dorkenwald et al., 2022; Zheng et al., 2018) and FANC (Phelps et al., 2021) datasets except for DNp03, for which the right VNC axon was used as it crosses the midline before descending on the contralateral side of the VNC.

#### EM mesh skeletonization

The raw morphological data obtained from the FAFB and FANC datasets provided surface reconstructions of neuron morphologies in the form of meshes, which are collections of points in space that connect to form polygons. To process the meshes for simulation in NEURON, the surface reconstructions were converted into skeletons containing radius information. A semi-automated pipeline was developed to reduce a complex mesh surface reconstruction into a simple skeleton model for any given neuron. This was achieved using the wavefront automated skeletonization algorithm from the *skeletor* Python package (https://navis-org.github.io/skeletor/). The algorithm casts waves along a mesh, connecting vertices into rings throughout the structure. The rings are then collapsed into single points centered within the original neuronal mesh while maintaining a calculated radius derived from the collapsed rings. By applying this algorithm, a morphologically accurate skeleton was extracted from EM-reconstructed neuronal meshes. All meshes were manually edited through *skeletor’s* pre-fix method to remove any disconnected fragments. For the wavefront algorithm, default parameters of waves = 1, and step size = 1 were used. Post skeletonization, *skeletor’s* post-clean method was used to remove any branches that ran parallel to their parent branch and move nodes outside the mesh back inside.

The FAFB and FANC neuronal mesh reconstructions of DNs are highly detailed as a result of their EM-level resolution and contain large numbers of vertices and polygons, which complicated the skeletonization process. Like most skeletonization algorithms, when the wavefront algorithm processed a large number of vertices, it often generated an excessive number of nodes, particularly for thicker neuronal processes, which led to excess branching. This algorithm, like others, also failed to consolidate the soma into a single node. Instead, it created a cluster of connected nodes branching off from each other. Skeleton models generated through this automated process were therefore manually proofread before being utilized in any simulations. Although this algorithm required manual proofreading, it faithfully maintained biological radius information unlike other skeletonization algorithms and substantially decreased the model generation time in comparison to traditional hand tracing.

For proofreading, MeshLab (Cignoni et al., 2008) was utilized to visualize the surface reconstruction and neuTube (Feng et al., 2015) was used for manual editing of the skeleton models. Somata were manually generated for each DN skeleton model using the average radius of the original soma cluster and corroborated with confocal microscopy images. Erroneous spinal branches and zero-radius nodes along the dendrites and axon were removed. Proofread skeleton models and the corresponding original mesh reconstructions were then overlaid to visually compare the final models. The total surface area of the final models was compared to the original surface area of the EM meshes to confirm the accuracy of the radius information. The DNp01 model required additional manual adjustments in the regions of the axon, soma, and tether (the process that connects the soma with the rest of the neuron) to more accurately represent the surface area, as the thick processes of DNp01 caused the algorithm to generate multiple nodes within the mesh, resulting in an underestimation of the radius of those nodes. Using neuTube the individual nodes throughout the skeleton models were labeled to represent their respective morphological elements to later facilitate efficient categorical access of sections within the simulation environment.

#### Axon estimation

The FAFB dataset is limited to neuron morphologies within the brain, however, DNs extend their axons into the VNC of the fly. To account for the missing axon sections in the models, the FANC (Phelps et al., 2021) dataset was accessed for morphological data on DN axons. To minimize the complexity of the model, the VNC mesh retrieved from the FANC dataset was skeletonized as outlined above. The axon skeleton was then reduced into a simplified equivalent cylinder that retained the overall surface area of the detailed skeleton and length estimated from published data (Namiki et al., 2018). The equivalent cylinder dimensions were then used to define an axon section that was appended to the FAFB-derived skeleton once it was loaded into the NEURON simulation environment.

**Table 1.**
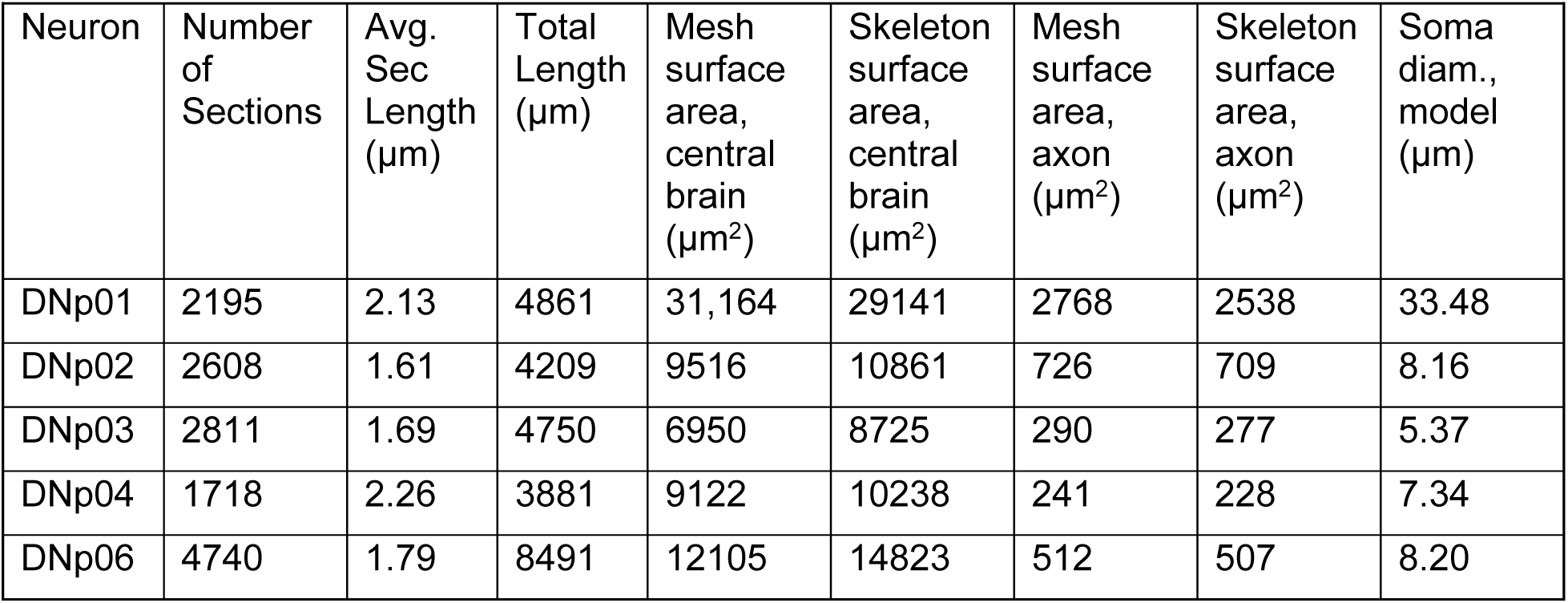
Model Morphological Parameters.

### Computational modeling and simulations

#### Model initialization

The model was initialized within the NEURON (Hines and Carnevale, 2001) simulation environment by loading the skeletonized FAFB morphology as an .swc file and then appending the artificial axon to the corresponding DN model. Optimized passive property values were then assigned to each section, with all sections having the same passive property set. The length constant λ was calculated for each section using its geometric and passive properties, and any section > 0.1 λ in length was discretized such that no segment within that section is longer than 0.1 λ. Next, all synapses were initialized at their appropriate locations by referencing the created synapse maps (see *Modeling of VPN-DN Synapses*). All simulations involving current injections or activation patterns of synapses were conducted using models with optimized passive properties and mapped synapses.

#### Optimization of passive properties

Optimal passive properties [capacitance (*C_m_*), leak conductance (*g_leak_*), axial resistivity (*R_a_*), and reversal potential (*E_rev_*)] were found by following a protocol established in previously published models of *Drosophila* neurons where model responses were fit to small hyperpolarizing current injections (Gouwens and Wilson, 2009; Günay et al., 2015; Hafez et al., 2023; Liu et al., 2022). Established experimental ranges of published model parameters were used to constrain biologically plausible ranges for the axial resistivity, *R_a_* (30-400 Ω*cm), and capacitance, *C_m_* (0.6-2.6 uF/cm^2^), for DN models (Borst and Haag, 1996; Gouwens and Wilson, 2009). The electrode was modeled as a simple cylindrical section (Gouwens and Wilson, 2009; Günay et al., 2015) with a length of 10 µm, a diameter of 1 µm, and the following passive properties: *R_a_* = 235.6 Ω*cm (emulating a total electrode resistance, of 30 MΩ), *g_leak_* = 3.798e-4 (emulating a total electrode seal resistance of 8 GΩ), and C_m_ = 6.4 µF/cm^2^ (emulating a total electrode capacitance of 2 pF).

The biophysical parameters for the model DNp01 (*g_leak_* = 4.35e-4 S/cm^2^, *E_rev_* = −66.63 mV, *R_a_* = 212 Ω*cm, *C_m_*= 0.7 uF/cm^2^) and DNp03 (*g_leak_* = 3.17e-4 S/cm^2^, *E_rev_* = −61.15 mV, *R_a_* = 50 Ω*cm, *C_m_*= 0.8 uF/cm^2^) were determined by manually adjusting passive property values within physiological ranges (Borst and Haag, 1996; Gouwens and Wilson, 2009) and minimizing an error function defined by the squared difference between the averaged experimental traces and the simulated trace.

#### Modeling of VPN-DN synapses

Synaptic inputs for each DN were retrieved from the FAFB datasets. For each DN, a list of synapses, along with coordinate information, presynaptic neuron ID, and a confidence metric (cleft_score) for each synapse was obtained. Individual synaptic inputs were labeled to correspond to their VPN cell type identity by comparing presynaptic neuron IDs to the published codex (version 630; https://codex.flywire.ai/app/search) (Dorkenwald et al., 2022). Synapses were filtered for quality by thresholding with a “cleft_score” of 30. The flywire default cleft_score of 50 was lowered to avoid underreporting synapse counts for the LPLC2 and LC4 inputs onto DNp01 when compared to known hand-traced counts from other published work (Ache et al., 2019). With this adjusted threshold, VPN-DN synapse counts were similar to those from the hemibrain EM dataset (Scheffer et al., 2020).

Synapses were mapped onto the morphology of their respective DN by calculating the Euclidean distance between the synaptic coordinates and every single coordinate point belonging to the skeleton. The synapse was then assigned to the section parenting the skeletal point closest to the synapse. The resulting synapse map was then saved for future reference. The resulting synapse maps and .swc morphologies for each model were imported into the Trees Toolbox (Cuntz et al., 2010) in Matlab to generate dendrogram visualizations. All available synapses from the FAFB dataset were mapped onto the skeletons, though only synaptic inputs belonging to VPNs were labeled and considered for this study.

The VPN cell types investigated here have been shown to be cholinergic and their synapses were therefore modeled accordingly (Davis et al., 2020; Dorkenwald et al., 2022; Eckstein et al., 2020; Schlegel et al., 2023). All synaptic conductances were modeled as a sum of two exponentials (using NEURON’s exp2syn method) utilizing time constants from previously modeled cholinergic synapses, with a rise time of 0.2 ms and a decay time of 1.1 ms. (Liu et al., 2022; Tobin et al., 2017). The maximum conductance was set to *g_max_* = 0.27 nS, and the synaptic reversal potential was set to *E_syn_* = –10 mV, consistent with previously modeled and reported *Drosophila* cholinergic synapses (Gouwens and Wilson, 2009; McCarthy et al., 2011; Phelps et al., 2021; Tobin et al., 2017). Synapse activations were driven by using the NetCon method in NEURON to link a stimulus event at a predefined time to the synapses of interest.

### Synaptic spread calculations

The synapse spread, *s*, was calculated as the average distance between all synapse pairs within a group as:

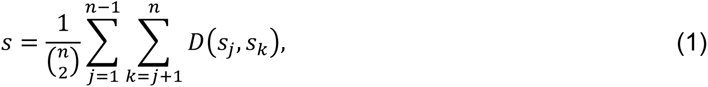

where *n* represents the number of synapses within a group, and *D(s_j_, s_k_*) represents the path distance along the DN between the *j*-th and *k*-th VPN synapse, as determined by NEURON’s distance function. The outer and inner sums iteratively cover all possible unique pairs of synapses, and the total sum is then divided by the total number of possible unique pairs, resulting in a final metric that represents the average distance between any two synapses within a group.

### Receptive field estimation

Individual VPNs were mapped to a location in the visual field based on their dendritic projections by following a previously established method (Dombrovski et al., 2023; Morimoto et al., 2020). LC4 and LC6 anatomical receptive fields were previously mapped using manually traced and proofread neurons in the FAFB catmaid interface (Dombrovski et al., 2023; Morimoto et al., 2020). The same FAFB EM dataset neurons were analyzed here but traced through the flywire.ai initiative which used artificial intelligence (AI) to trace neuron morphologies and independent manual proofreading through crowd-sourced initiatives (Dorkenwald et al., 2022). Both approaches yielded comparable results which also confirmed that possible discrepancies in manual proofreading did not affect receptive field estimations.

For each VPN population, the dendritic arbors were mapped within the lobula and a second-order polynomial was fit through those dendrites to create a quadratic plane, estimating the curvature of the lobula (Morimoto et al., 2020). An exception was made for LPLC4 because the dendritic arbors of the LPLC4 population showed a bias towards the anterior portion of the lobula, which caused the quadratic plane and subsequently the estimated receptive fields to be distorted. The LPLC1 plane was therefore instead utilized for LPLC4 because the two populations share similar dendritic arborizations in layers 2-4, with the only key difference being that LPLC1 also arborizes in layer 5b while the LPLC4 population also arborizes in layer 6 of the lobula (Wu et al., 2016).

Polygons were generated that represent the estimated field of view of individual neurons in the population by identifying where the dendritic branches of each neuron intersect the quadratic plane. From those same dendritic arbors, a centroid point representing the center of the receptive field of that given neuron was calculated. The VPN polygons were then mapped with their respective centroid points onto eye coordinates representing the fly’s visual field. Previously reconstructed Tm5 neurons were utilized as known reference points that correspond to the center of the eye and dorsal position on the central meridian (Morimoto et al., 2020).

### Statistical analysis

All statistical analyses were carried out using the SciPy (Virtanen et al., 2020) and scikit-posthocs (Terpilowski, 2019) packages in Python. Normality of all sample datasets was tested using the Shapiro-Wilk test (**Figures 5** and **12**). Linear regression analysis was used to test for retinotopy (**Figure 4D-E and Figure 4—figure supplement 1**). For all comparisons where a non-normal distribution was found, a non-parametric Mann–Whitney U test was used to compare the groups (**Figure 5B**). For comparisons in the nearest neighbor analysis (**Figure 5C**), a one sample t-test was used to compare the means of shuffled trials to the original dataset. Comparisons across multiple groups (**Figure 12** and **Figure 12—figure supplement 1**) were performed using Kruskal-Wallis tests with a p-value <0.05 required for significance. Significant results were followed up with a Dunn’s post-hoc test with a Bonferroni adjustment of p-values (Dunn, 1964; Glantz, 2012).

## ACKNOWLEDGEMENTS

We thank Arthur Zhao for help with the receptive field mapping, James M. Jeanne for help with the creation of dendrograms, and Thomas A. Ravenscroft for providing us with para-GFSTF tools for SIZ labeling. Confocal experiments were conducted at Drexel University’s Cell Imaging Center, RRID:SCR_022689

This study was supported in part by the National Institutes of Health (NINDS R01NS118562 to J.A. and C.R.v.R.), and the National Science Foundation (grant no. IOS-1921065 to C.R.v.R.).

## AUTHOR CONTRIBUTIONS

Conceptualization: C.R.v.R., J.A.

Methodology: A.M-S., A.N.V., C.R.v.R, J.A.

Software: A.M-S., A.N.V.

Investigation: A.M-S., A.N.V., H.J., B.W.H., C.R.v.R, J.A.

Resources: C.R.v.R, J.A.

Data Curation: A.M-S., A.N.V.

Writing – Original Draft: A.M-S., A.N.V., J.A.

Writing – Review & Editing: A.M-S., A.N.V., H.J., B.W.H., C.R.v.R, J.A.

Visualization: A.M-S., A.N.V., B.W.H., J.A.

Supervision: C.R.v.R, J.A.

Funding Acquisition: C.R.v.R, J.A.

## DECLARATION OF INTERESTS

The authors declare no competing interests.

## SUPPLEMENTAL FIGURES

**Figure 3—figure supplement 1.**
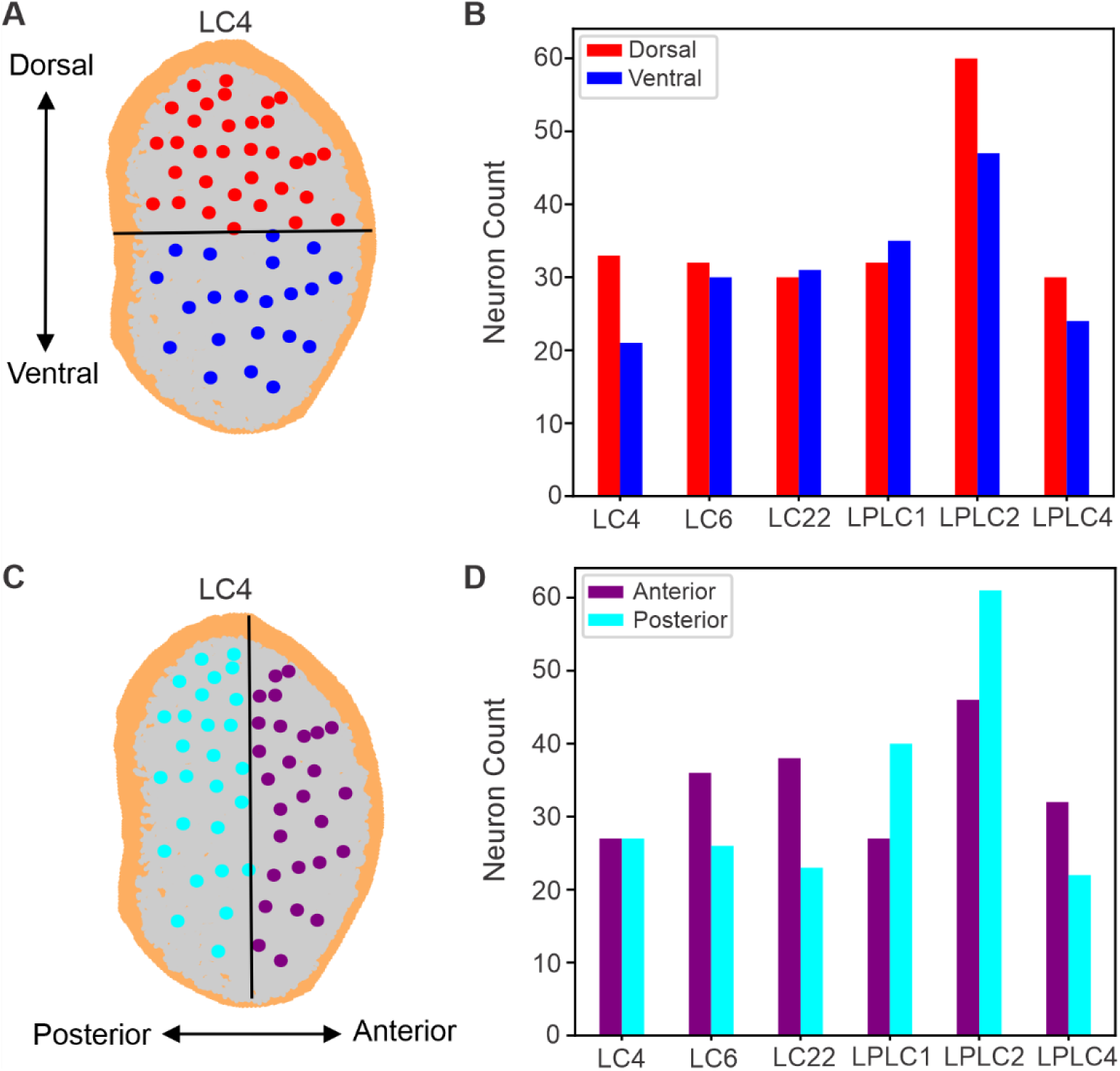
VPN centroids tile the lobula with biases across the two main axes. (**A**) Example LC4 centroid (dorsal: red, ventral: blue) and their dendrite (gray) projections onto the lobula. The black line splits the lobula in half along the dorso-ventral axis. (**B**) Number of VPN centroids in the dorsal and ventral lobula hemisphere. Some VPN populations show biases. (**C, D**) Same as in (A, B) but for the posterior (cyan) and anterior (purple) lobula hemispheres.

**Figure 4—figure supplement 1.**
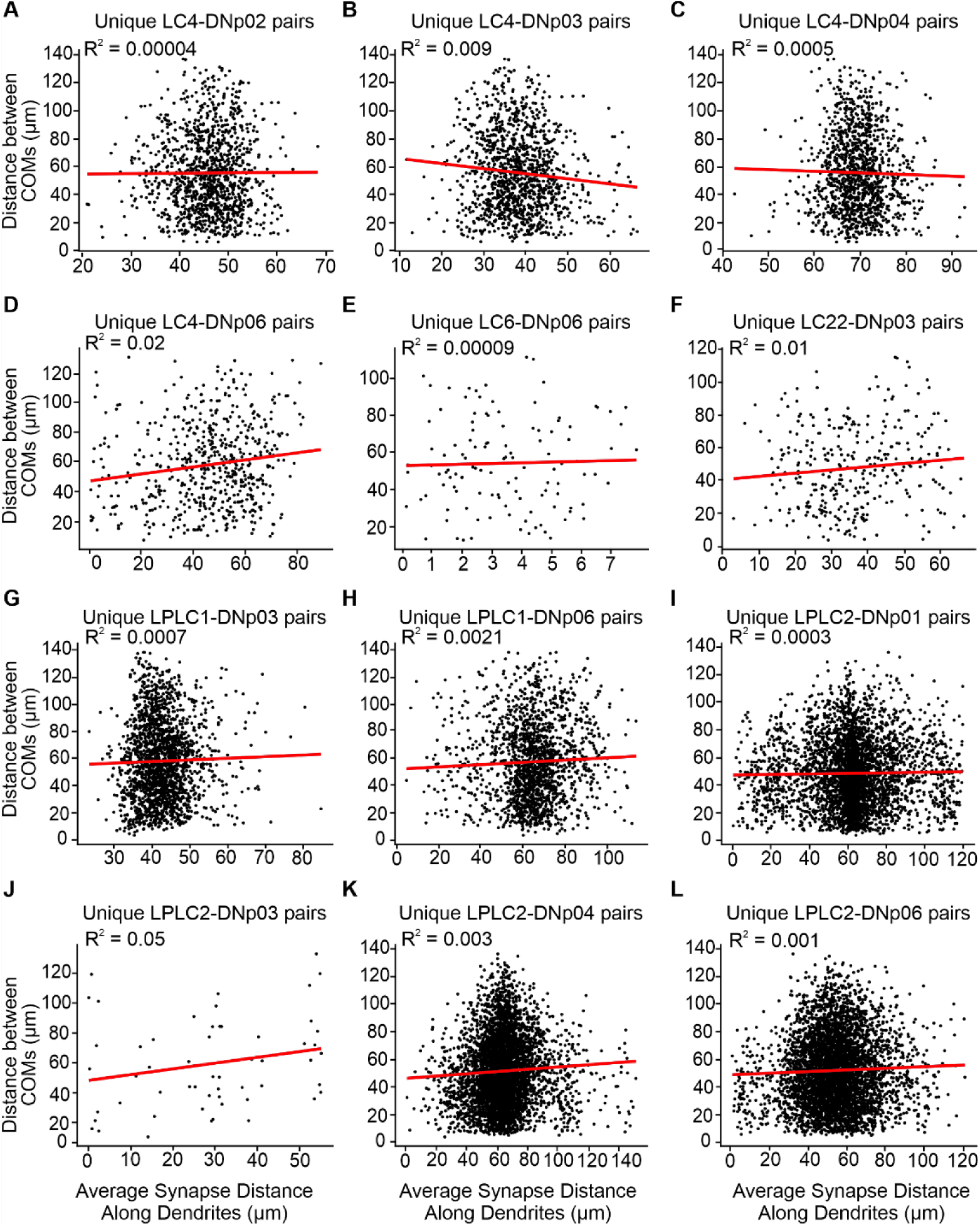
Lack of retinotopic synapse organization in all VPN-DN pairs. Linear regression line in red. **(A-L)** (**A**) The distance between the centroids of all unique pairs of LC4 presynaptic to DNp02 is not correlated with the average synapse distance for that given pair of LC4 neurons. (**B**) Same as (A) but for LC4-DNp03. (**C**) Same as (A) but for LC4-DNp04 (**D**) Same as (A) but for LC4-DNp06 (**E**) Same as (A) but for LC6-DNp06 (**F**) Same as (A) but for LC22-DNp03 (**G**) Same as (A) but for LPLC1-DNp03 (**H**) Same as (A) but for LPLC1-DNp06 (**I**) Same as (A) but for LPLC2-DNp01 (**J**) Same as (A) but for LPLC2-DNp03 (**K**) Same as (A) but for LPLC2-DNp04 (**L**) Same as (A) but for LPLC2-DNp06

**Figure 8—figure supplement 1.**
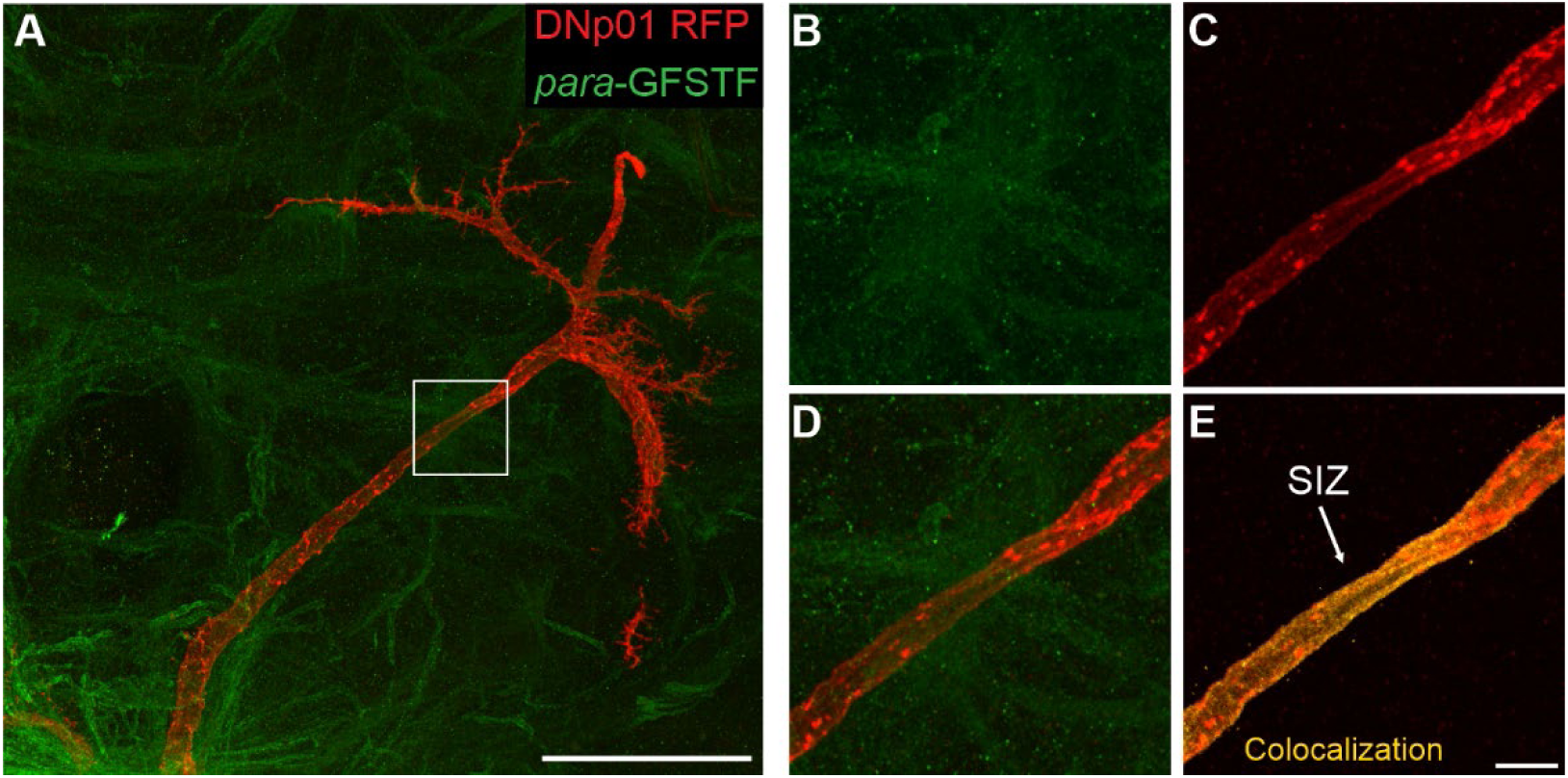
The SIZ is located downstream of the tether in DNp01. (**A**) Maximum intensity projection of RFP labeled DNp01 and GFP labeled para (genotype: *para-GFSTF, DNp01-split-GAL4, UAS-RFP).* Scale bar: 50 µm. Crosses were set at 22°C. (**B-E**) Zoomed-in view of (A) showing (**B**) the individual para and (**C**) DNp01 channels, (**D**) their overlay, and (**E**) para colocalization on DNp01 Scale bar: 5 µm.

**Figure 8—figure supplement 2.**
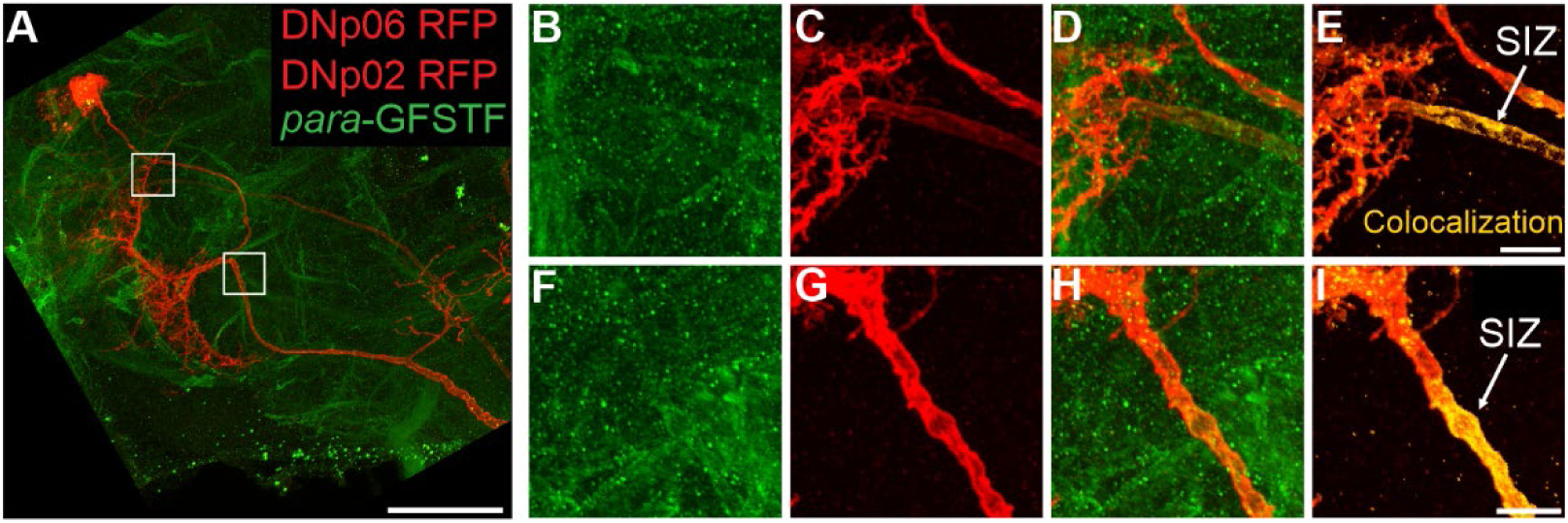
The SIZ is located downstream of the tether in DNp02 and DNp06. (**A**) Maximum intensity projection of RFP labeled DNp06 and DNp02 and GFP labeled para (genotype: *para-GFSTF, DNp06-split-GAL4, UAS-RFP*). Scale bar: 50 µm. Crosses were set at 18°C. (**B-E**) Zoomed-in view of top white box in (A) showing (**B**) the individual para and (**C**) DNp06/DNp02 channels, (**D**) their overlay, and (**E**) para colocalization on DNp06. Scale bar: 5 µm. (**F-I**) Zoomed-in view of bottom white box in (A) showing (**F**) the individual para and (**G**) DNp06/DNp02 channels, (**H**) their overlay, and (**I**) para colocalization on DNp02. Scale bar: 5 µm.

**Figure 8—figure supplement 3.**
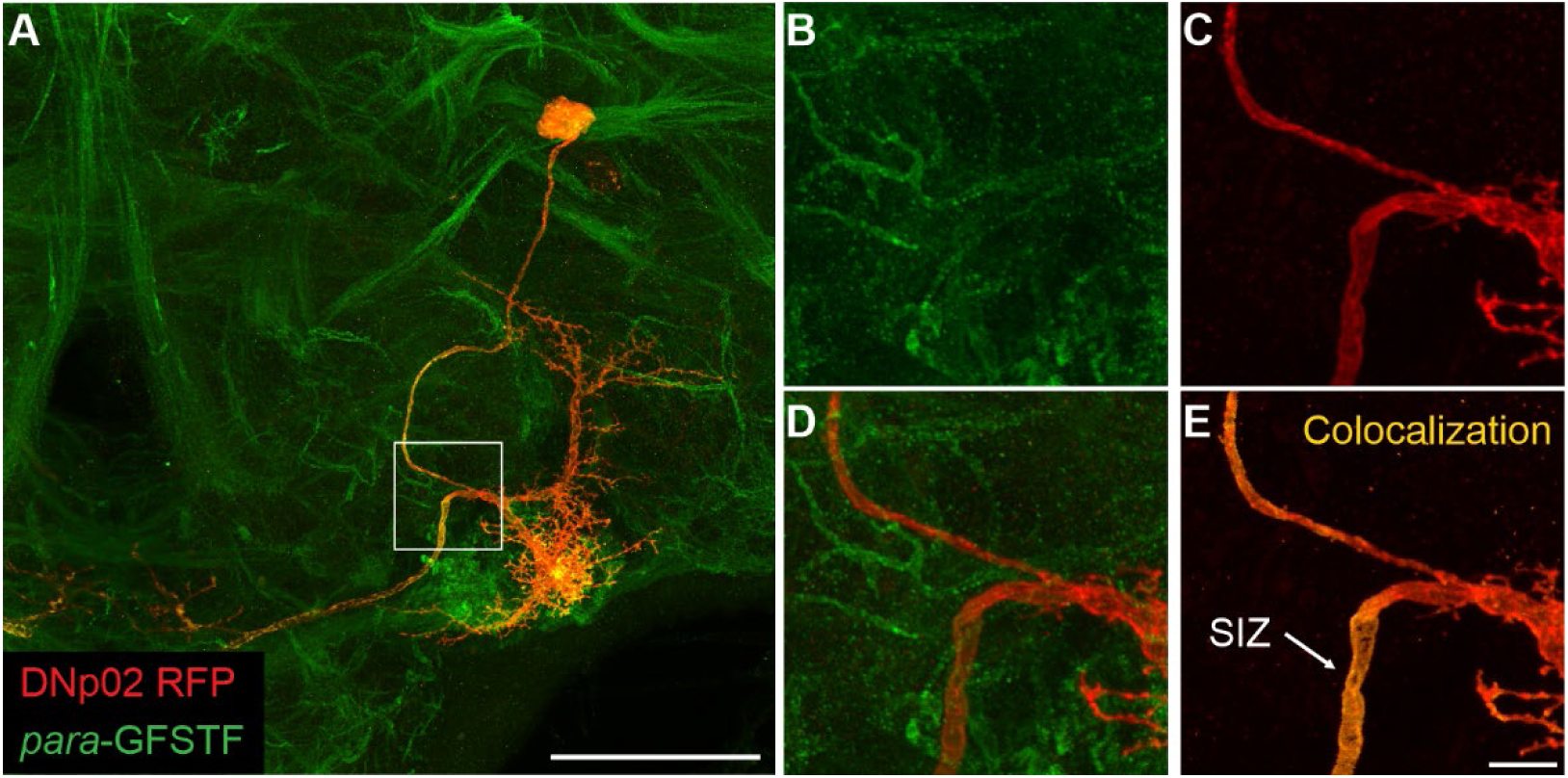
The SIZ is located downstream of the tether in DNp02. (**A**) Maximum intensity projection of RFP labeled DNp02 and GFP labeled para (genotype: *para-GFSTF, DNp02-split-GAL4, UAS-RFP).* Scale bar: 50 µm. Crosses were set at 18°C. (**B-E**) Zoomed-in view of (A) showing (**B**) the individual para and (**C**) DNp02 channels, (**D**) their overlay, and (**E**) para colocalization on DNp02 Scale bar: 5 µm.

**Figure 12—figure supplement 1:**
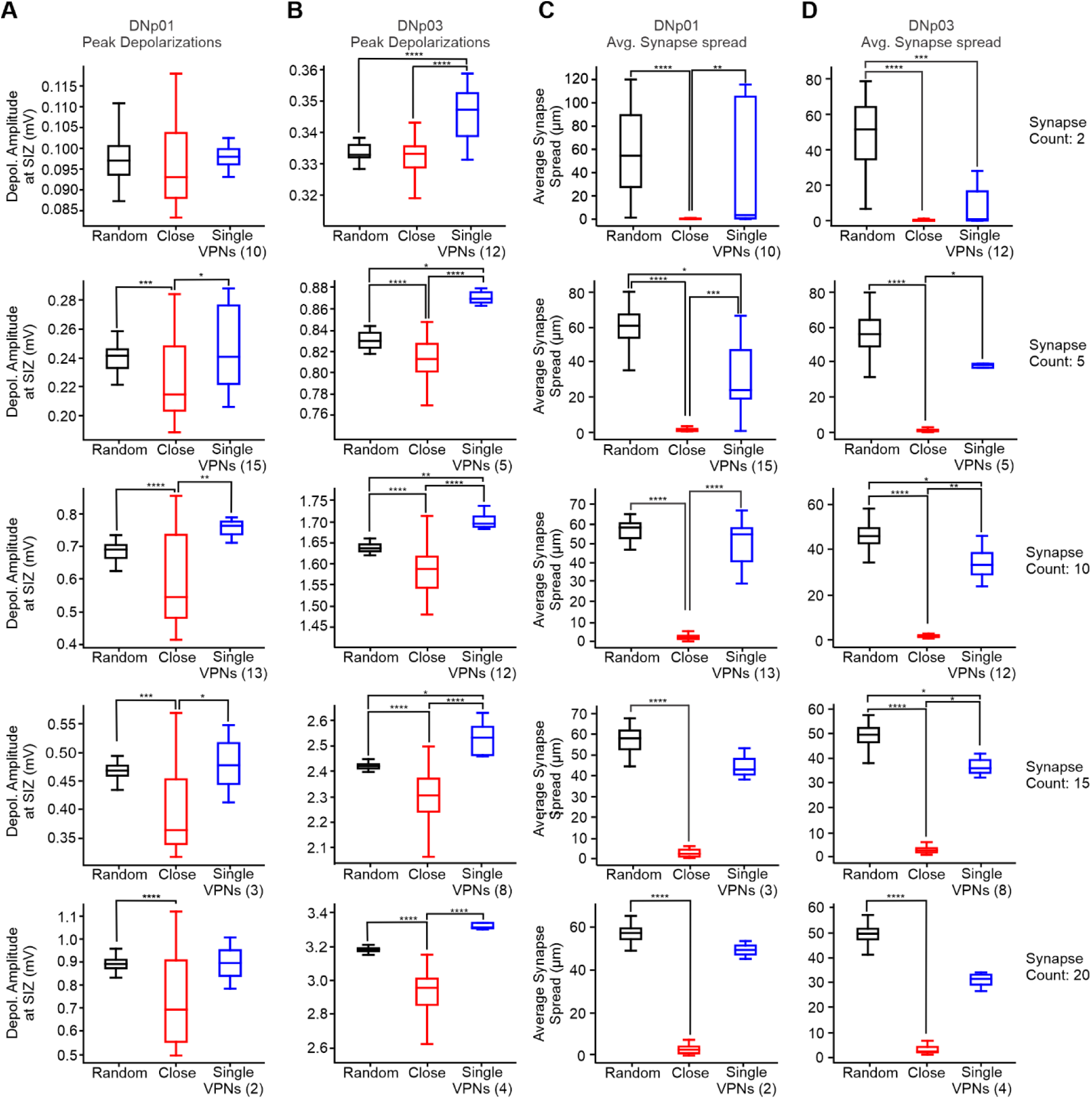
VPNs distribute their synapses to achieve efficient composite EPSP amplitudes at the SIZ. **(A, B)** Composite EPSP amplitudes at the SIZ in response to simultaneous activation of varying numbers of synapses in DNp01 (A) and DNp03 (B). **(C,D)** Average synapse spread of randomly distributed synapses, closely clustered and single VPNs at varying synapse counts in DNp01 (A) and DNp03 (B).

## Notes

### Competing Interest Statement

The authors have declared no competing interest.

